# Oscillation-driven memory encoding, maintenance and recall in an entorhinal-hippocampal circuit model

**DOI:** 10.1101/804062

**Authors:** Tomoki Kurikawa, Kenji Mizuseki, Tomoki Fukai

## Abstract

During the execution of working memory tasks, task-relevant information is processed by local circuits across multiple brain regions. How this multi-area computation is conducted by the brain remains largely unknown. To explore such mechanisms in spatial working memory, we constructed a neural network model involving parvalbumin-positive, somatostatin-positive and vasoactive intestinal polypeptide-positive interneurons in the hippocampal CA1 and the superficial and deep layers of medial entorhinal cortex (MEC). Our model is based on a hypothesis that cholinergic modulations differently regulate information flows across CA1 and MEC at memory encoding, maintenance and recall during delayed nonmatching-to-place tasks. In the model, theta oscillation coordinates the proper timing of interactions between these regions. Furthermore, the model predicts that MEC is engaged in decoding as well as encoding spatial memory, which we confirmed by experimental data analysis. Thus, our model accounts for the neurobiological characteristics of the cross-area information routing underlying working memory tasks.

## Introduction

Spatial navigation is a fundamental cognitive function that requires the processing of spatial memory by the hippocampus and entorhinal cortex. During a spatial navigation task, spatial information relevant to the task has to be encoded into, maintained in and recalled from spatial working memory at behaviorally adequate times. How these operations are coordinated by the cortico-hippocampal neural circuits during a spatial working memory task has yet to be explored.

A spatial working memory task is processed by several cortical areas such as the medial prefrontal cortex (mPFC) (Benchenane et al., 2010; Jones & Wilson, 2005; Spellman et al., 2015), medial entorhinal cortex (MEC)(Suh, Rivest, Nakashiba, Tominaga, & Tonegawa, 2011; Yamamoto, Suh, Takeuchi, & Tonegawa, 2014) and the hippocampal area CA1 (Benchenane et al., 2010). These anatomically connected areas (Eichenbaum, 2017; Swanson & Cowan, 1977; Witter, Wouterlood, Naber, & Van Haeften, 2000) are thought to mutually communicate information necessary to accomplish the task. Importantly, the degree of functional importance of different inter-area connections varies during the task. This is indicated by the fact that the impairment of these connections at different behavioral phases differentially influences task performance (Spellman et al., 2015; Suh et al., 2011; Yamamoto et al., 2014). For instance, in a delayed nonmatching to place task (DNMP), the maintenance of spatial memory during a delay period does not require synaptic connections from the layer 3 of the MEC (MECIII) to CA1, but these connections are necessary for memory recall (Yamamoto et al., 2014). Connections from CA1 to the mPFC play a crucial role in memory encoding but not in memory recall (Spellman et al., 2015). These results indicate that information flows via the hippocampal circuit are not static but are dynamically regulated depending on the behavioral demands.

Dynamic information routing across multiple areas is thought to reflect in coherence in neuronal activity between different areas (Spellman et al., 2015; Yamamoto et al., 2014), which leads to the hypothesis called “communication through coherence”(Fries, 2015). Many theoretical (Akam & Kullmann, 2010; Buehlmann & Deco, 2010; Palmigiano, Geisel, Wolf, & Battaglia, 2017; Vogels & Abbott, 2005; Yang, Murray, & Wang, 2016) and experimental (Letzkus, Wolff, & Lüthi, 2015; Womelsdorf, Valiante, Sahin, Miller, & Tiesinga, 2014) studies have explored the gating functions for this dynamic processing. However, how the computations installed at multiple cortical areas are integrated to execute a spatial working memory task including different cognitive stages (encoding, maintenance, and decoding) remains largely unclear.

Here, we elucidated the underlying mechanisms of multi-area dynamic information processing during DNMP tasks. We hypothesize that acetylcholine (ACh) controls spatial information flows in the entorhinal-hippocampal circuit according to different cognitive demands. Indeed, ACh is involved in diverse cognitive functions (Hasselmo & Sarter, 2010; Parikh, Kozak, Martinez, & Sarter, 2007) including fear conditioning (Letzkus et al., 2011; Pi et al., 2013), sensory discrimination (Hangya, Ranade, Lorenc, & Kepecs, 2015; Pinto et al., 2013), associative memory (Sabec, Wonnacott, Warburton, & Bashir, 2018), and spatial (Croxson, Kyriazis, & Baxter, 2011; Okada, Nishizawa, Kobayashi, Sakata, & Kobayashi, 2015) and non-spatial working memory tasks (Furey, Pietrini, & Haxby, 2000; Hasselmo, 2006; McGaughy, Koene, Eichenbaum, & Hasselmo, 2005).

To test the hypothesis, we constructed a biologically plausible model of the entorhinal-hippocampal circuit consisting of MECIII, CA1 and MEC layer V (MECV) with ACh projections from the medium septum and numerically simulated a DNMP task on a T maze. We show that the cholinergic modulation of disinhibitory circuit in CA1 and a calcium-dependent cation current in MEC is crucial for coordinating the encoding, maintenance and retrieval modes of the MECIII-CA1-MECV circuit. The model successfully reproduces theta phase preferences in various types of CA1 (Klausberger & Somogyi, 2008) and MEC (Mizuseki, Sirota, Pastalkova, & Buzsáki, 2009) neurons. Further, we demonstrate whether the inactivation of MECIII-to-CA1 input may impair performance in DNMP tasks depends on the timing of inactivation, as was shown in experiment (Yamamoto et al., 2014).

Our model also predicts that the same MECIII neurons encoding spatial information retrieve this information later, which we confirm by analyzing experimental data of single-cell and population-level activities. According to a widely accepted view, CA3-to-CA1 input is responsible for retrieving spatial memory (Fernández-Ruiz et al., 2017; S. J. Middleton & McHugh, 2016). However, a few studies suggested that MECIII-to-CA1 input is engaged in the recall of spatial memory (Suh et al., 2011; Yamamoto et al., 2014), and our model supports the latter view.

## Results

### Hippocampus-entorhinal cortex circuit model

To clarify the circuit mechanisms to control flexibly spatial information in the hippocampus and MEC, we built an inter-areal cortical network model (Figure 1A, see Supplemental materials for details). The network comprises three main areas CA1, MEC layer 3 (MECIII) and layer 5 (MECV), and includes additional areas CA3, MEC layer 2 (MECII) and the medial septum (MS) as external inputs. These external inputs oscillate at theta frequency (10Hz) and entrain the main circuit to theta-frequency oscillation. CA3 neurons encode the current location of model rat and transfer the location information to CA1. All main areas have excitatory (E) and parvalbumin (PV)-positive interneurons. In addition to these neurons, the model CA1 has somatostatin (SOM)-positive oriens-lacunosum moleculare (OLM) and vasoactive intestinal polypeptide (VIP) neurons. We built synaptic connections in our model based on anatomical observations (Gonzalez-Sulser et al., 2014; Unal, Joshi, Viney, Kis, & Somogyi, 2015; Witter et al., 2000). In addition, we assumed that E neurons in MECII project to PV neurons in MECIII, as previously suggested (Mizuseki et al., 2009).

**Figure 1.**
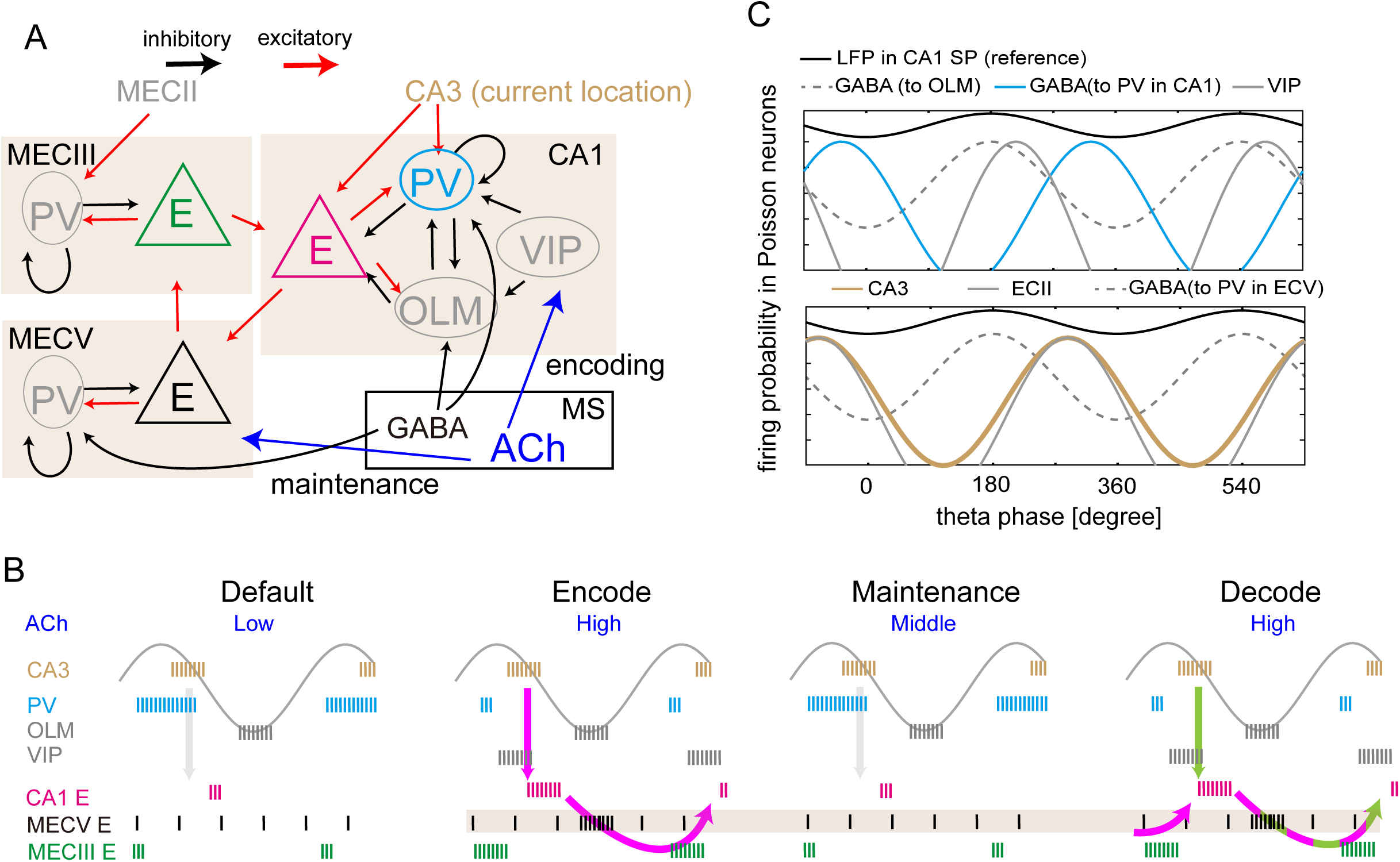
Entorhinal-hippocampal network model (A) Connectivity of the network model is shown. (B) Schematic image of the proposed mechanisms of spatial working memory. Vertical lines represent spikes of each neuron. Sinusoidal curve indicates theta oscillation in the CA1 stratum pyramidale (SP) layer. As described in C, this theta oscillation was used for a reference frame to measure the preferred phases of neuronal firing in our model. See the main text for details. (C) Theta-modulated firing probabilities of input neurons are shown during the sample-center run. The preferred firing phases were determined with respect to a reference theta oscillation (black) reported in the CA1 SP.

Acetylcholine (ACh) is known to modulate activity of VIP neurons (Albuquerque, Pereira, Alkondon, & Rogers, 2009) and the conductance of calcium-sensitive non-specific cation current (CAN) in MECV excitatory cells (Fransen, Alonso, & Hasselmo, 2002; Fransén, Tahvildari, Egorov, Hasselmo, & Alonso, 2006). The concentration of ACh ([ACh]) is generally thought to change in a diffusive and tonic manner on slow timescales of minutes or hours. However, recent studies have revealed that [ACh] undergoes rapid phasic changes on sub-second and second timescales (Parikh et al., 2007; Teles-Grilo Ruivo et al., 2017; H. Zhang, Lin, & Nicolelis, 2010). In this study, we assumed that [ACh] varies in correlation with cognitive demands, i.e., [ACh] is high, slightly lowered and again high during encoding, maintenance and recalling of working memory, respectively. In contrast, the default concentration of Ach is low (Figure 1B). In reality, these cholinergic modulations may be induced by MS (Newman, Gupta, Climer, Monaghan, & Hasselmo, 2012; Okada et al., 2015; H. Zhang et al., 2010), but cholinergic neurons in MS were not explicitly modeled in the present study. Under this model setting, we tested the hypothesis that ACh controls information flow across the entorhinal-hippocampal loop circuit in the different cognitive stages of a DNMP task. In particular, we proposed and explored the possibility that [Ach] regulates the disinhibition of CA1 E neurons and the calcium dynamics in MECV E neurons to enable the flexible processing of spatial working memory.

Before showing the details of our network model, we schematically explain the mechanisms of spatial working memory which we intend to propose in this study (Figure 1B). In the default stage (before encoding), [ACh] is low and activity of PV is high enough to inhibit activity of CA1 E neurons and the flow of information on the current location from CA3 to CA1 (shaded arrow in the panel of default stage). In the encoding stage, [ACh] is set higher and consequently VIP neurons are activated, which in turn inhibits CA1 PV neurons to enable the transfer of location information into CA1. Thus, the location information is stored in the CA1-MECV-MECIII loop circuit (magenta arrows in the panel of encode stage). Next, in the maintenance stage, [ACh] is at a middle level and PV neuron activity is again high, blocking the re-entry of location information into CA1. Despite this blockade, however, the spatial information is maintained by the activation of calcium-dependent cation current in MECV E neurons without spike generation. We assume that the current remains activated at the modest level of [ACh]. Finally, in the decoding stage, [ACh] is high again and information on the current location is loaded from CA3 to CA1 (green arrow). It is noted that information on the previous location maintained in MECV is also re-loaded from MECIII to CA1. Therefore, CA1 exhibits both current and previous position-encoding activities in the decoding stage.

The core circuits of the present network model consist of E, PV and OLM neurons, which were modeled as Hodgkin-Huxley-type conductance-based neurons according to previous models (S. Middleton et al., 2008; Rotstein, Oppermann, White, & Kopell, 2006; Wang & Buzsáki, 1996; Wulff et al., 2009). VIP neurons were modeled as Poisson firing neurons with a probability density of spikes and their outputs reflect the modulatory effect of ACh. We described CA3 E neurons projecting to CA1, MECII E neurons projecting to MECIII, and GABAergic neurons in MS projecting to CA1 and MECV as external Poisson spike trains. As shown in Figure 1C, the firing probabilities of these neurons were modulated at the theta-band frequency (10 Hz) to induce theta rhythmic activities in the core circuits. The relative preferred theta phases of the external inputs were chosen such that the preferred phases of various model neurons are consistent with experimental observations (Klausberger & Somogyi, 2008; Mizuseki et al., 2009). Furthermore, the relative phases between these inputs and theta oscillation in CA1 are experimentally known, from which we can define the theta phase of the local field potential (LFP) to be observed in the stratum pyramidale (SP) of CA1. This LFP oscillation was used as a reference to measure the degree of agreement between the preferred phases of model neurons and experimental observations. The preferred phases of GABAergic neurons in MS were dependent on their target neuron types (ECV PV, CA1 PV and CA1 OLM). In experiment, GABAergic neurons projecting to OLM and those projecting to PV in CA1 have different preferred phases (Borhegyi, 2004).

Given these settings of external theta-rhythmic sources, in a default stage the resultant firing of all neurons in CA1, MECIII and MECV showed theta-rhythmic patterns and their preferred phases are consistent with experimental observations (Figure S1): E and OLM neurons in CA1 showed preferred phases around the troughs of theta oscillation, whereas PV neurons around the peaks (Klausberger & Somogyi, 2008). In MECIII, E and PV neurons fired preferentially around the peaks and troughs of theta oscillation, respectively (Mizuseki et al., 2009). In MECV, PV neurons preferred the troughs, but E neurons did not show strong phase preferences (Mizuseki et al., 2009). Thus, the model neurons show biologically plausible theta-phase locking activity.

**Figure S1 (related to Figure 1).**
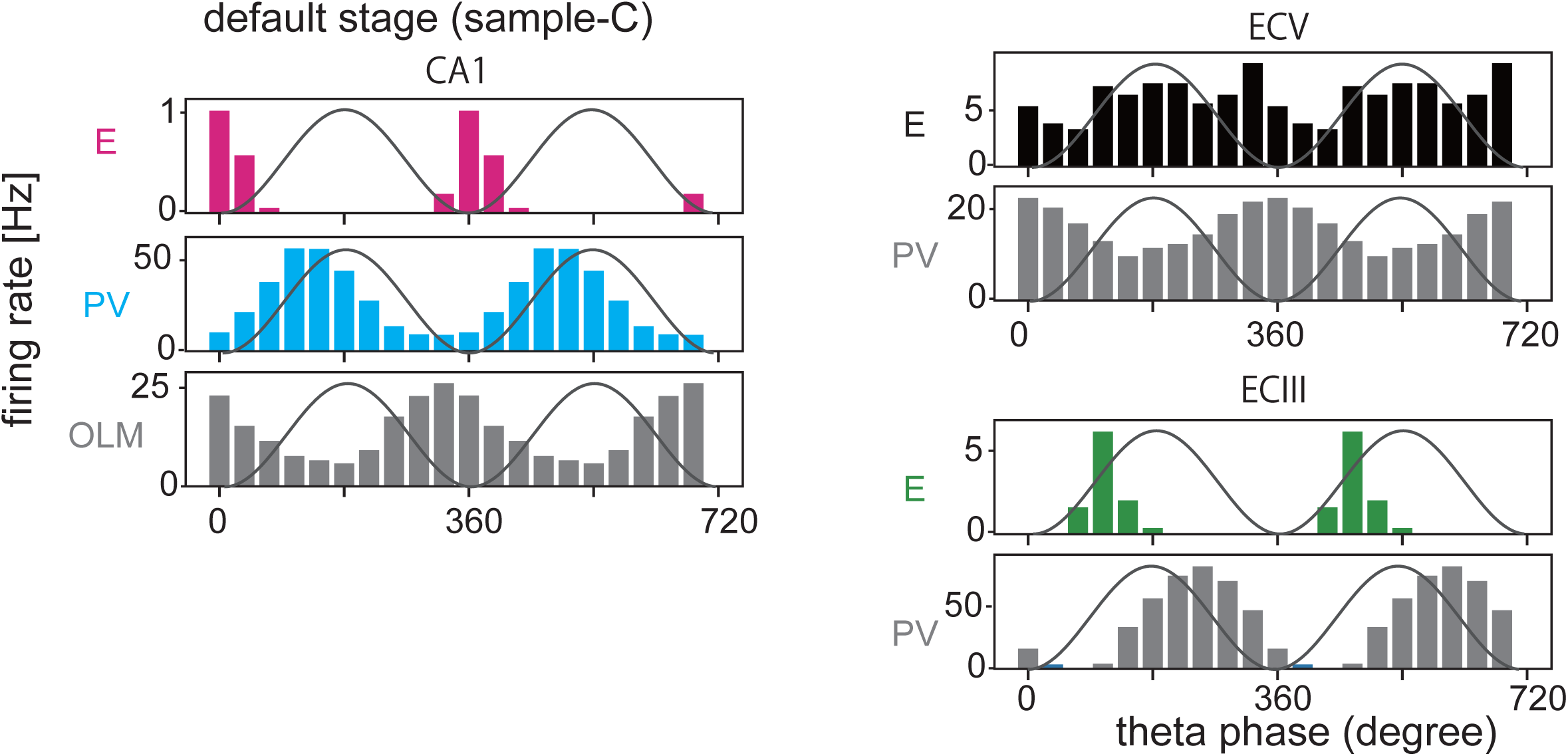
Activity of various types of neuron during task. Preferred phases of various neuron types during the sample-C period. The firing rates were calculated by numerical simulations for excitatory and inhibitory neurons in the entorhinal-hippocampal circuit. The reference theta oscillation presumed in the SP layer of CA1 is also shown (solid lines).

### Encoding and recalling of spatial information in the circuit model

The central question of this study is to clarify how spatial information is encoded, maintained and recalled in the entorhinal-hippocampal circuits. Before studying this problem, we, first, asked whether our model can replicate the task-related activities reported in previous experiments. In particular, we considered a DNMP task on a T-maze (Yamamoto et al., 2014). In this experiment, one arm of the T-maze was closed during a sample run and the mouse was forced to choose another arm. After a delay period, the mouse was set to a test run in which both arms were open and the mouse had to choose the arm opposite to the one chosen in the preceding sample run (that is, if the mouse chose the right arm in the sample run, it had to choose the left arm in the test run). For a successful test run, the mouse had to remember the previously chosen arm. This task requires at least two types of memory, namely, rule-based memory and spatial working memory. The former memory is thought to be encoded in the prefrontal cortex (Durstewitz, Vittoz, Floresco, & Seamans, 2010; Guise & Shapiro, 2017; Preston & Eichenbaum, 2013). However, in this study we did not model the prefrontal circuits and focused on the processing of spatial working memory in the entorhinal-hippocampal circuits.

Figure 2A shows the organization of sample and test runs in our model together with the connectivity patterns between the neural ensembles encoding different locations on the maze. We monitored the activity of sample runs along the center arm (sample-C) and left arm (sample-L), and that of test runs at the home position (delay) and center arm (test-C) up to the junction (decision point) of the T-maze. For the sake of simplicity, we implemented four subgroups L, R, C and H of place cells in CA3, each of which encoded the current position of the model mouse on the left, right, center arms and at the home position, respectively. For instance, neurons belonging to the subgroup L were given a higher firing probability when the model mouse traveled across the left arm (Figure 2B; also see “Neuron models” in Materials and methods). Accordingly, E neurons in CA1 were also divided into four subgroups L, R, C, and H, each of which was strongly projected to by the corresponding subgroup in CA3. In contrast, MECIII and MECV had two subgroups denoted as L and R and these subgroups were assumed to form closed loop circuits with the corresponding CA1 subgroups. In the present study, C and H subgroups are not necessary in MEC and we omitted these parts for the sake of simplicity. The other positions on the maze that are not shown in Figure 2A were not modeled.

**Figure 2.**
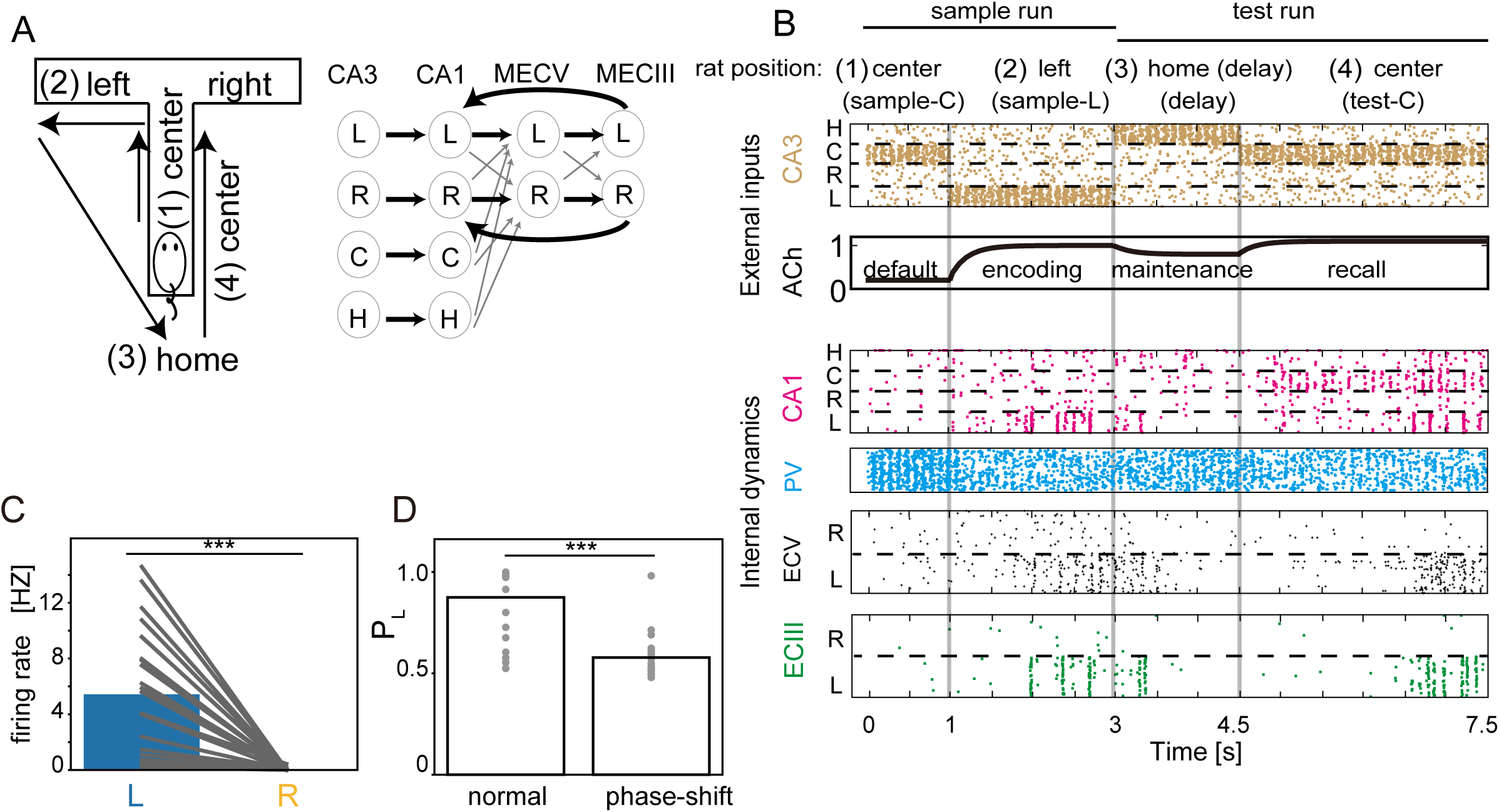
Performance of the model for a DNMP task (A) Left; Schematic illustrations of task periods during the sample and test trials of the DNMP T-maze task: (1) center in the sample trials (sample-C), (2) left in the sample trials (sample-L), (3) home (delay), (4) center in the test trials (test-C) periods. Right; Synaptic connectivity is shown between the neuron subgroups encoding the specific locations of the maze (left, right, center arms and home position). Connections (bold) are stronger within the loop circuit of MECV, MECIII and CA1 than other modest connections (solid). (B) Raster plots of E neurons in different cortical areas are shown together with the time evolution of ACh concentration. (C) The average firing rates of MECIII L and R subgroups were calculated for the test-C period in 25 runs of simulations (black lines, five different networks with five different initial conditions). Unless otherwise stated, average firing rates were evaluated in a similar fashion throughout this study. (D) Probability of left choice (P_L) and average of P_L was calculated in the normal and phase-shift conditions. Gray dots show P_L of different networks for different initial conditions. Chance level is 0.5.

Below, without loss of generality, we consider the case that the mouse chooses the left arm in every sample trial. Accordingly, we define task performance as the probability of Left choice, *P*_*L*_, which is estimated as 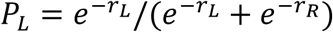, where *r*_*L*_ and *r*_*R*_ are the spike rates of the L and R subgroups, respectively, during the test-C period. Note that *P*_*L*_ can be written as a function of *r*_*L*_−*r*_*R*_. The larger the difference in the spike rates, the more robust the memory encoding.

In experiment, coherence between MECIII and CA1 increased in the high-gamma band (60-120Hz) as the animal approached the junction point on the T-maze (Yamamoto et al., 2014). The result indicates that high-gamma oscillation plays an active role in the decision making, presumably in reading out stored locations from working memory. However, the task performance of the animal was also highly correlated with theta-phase-locked firing in MECIII. In addition, as shown later, our model successfully performs working memory tasks without gamma oscillation. We speculate that high-gamma oscillation nested in theta oscillation may either help the downstream areas (responsible for the generation of behavioral outputs) to correctly read out the stored locations from the entorhinal-hippocampal circuit or mediate top-down signals to shift the status of the entorhinal-hippocampal circuit from the maintenance mode to the readout mode. In this study, we focus on spatial information processing within CA1 and MEC, and will not model any process arising outside of the entorhinal-hippocampal circuit.

### Implications of preferred theta phases in coordinating activities in the CA1-MECV-MECIII loop circuit

In Figure 2B, we show activities of E neurons in CA3, CA1, MECV and MECIII together with [ACh] during (1) sample-C, (2) sample-L, (3) home (delay) and (4) test-C runs. Depending on the mouse’s position, the corresponding subgroup was activated in CA3 according to the given firing probability. During the sample-L run, [ACh] was set to increase, which disinhibited CA1 PV neurons and accordingly strongly activated the CA1-MECV-MECIII loop circuit of the L subgroups to encode a choice memory in MECIII. Then, in the delay period, [ACh] was set to decrease slightly, which strongly suppressed neural activities in all L subgroups including the CA1 subgroup L. We note that a similar suppression arose in experiment as if spatial memory had not been maintained during delay periods (Yamamoto et al., 2014). During the test-C run, the CA1 subgroup C was activated driven by the CA3 subgroup C. Importantly, as the model mouse approached to the decision point, the subgroups L were gradually reactivated in the loop circuit to retrieve the memory of the previous choice. This activation-suppression-reactivation pattern is clearly seen in the firing rate of MECIII neurons. Average firing rates during the test-C period were significantly different (p=3.071×10^−6^, *t* test on two related samples) between the subgroups L and R in MECIII (Figure 2C), implying that the network model successfully recalled the stored spatial memory.

The successful encoding of memory required theta-phase-locking of neural firing along the CA1-MECV-MECIII loop circuit. As mentioned previously, the theta phases of external sources (i.e., CA3, MECII and MS) entrain neurons in these areas in theta-phase-locked firing with the preferred phases that are consistent with experimental observations. In the normal situation, MECIII PV neurons are activated by input from MECII at the troughs of theta oscillation and consequently MECIII E neurons tend to fire at the peaks. Then, CA1 E neurons are strongly activated in a non-linear manner(Bittner et al., 2015; Takahashi & Magee, 2009) by near-coincident inputs from MECIII and CA3: spikes from CA3 arriving at CA1 just after spikes from MECIII activate CA1 E neurons much stronger than those from CA3 arriving at otherwise timing (see Materials and methods).

The theta phases of inputs from the different sources were tuned such that they are consistent with experimental observations (Figure 1C). Is this coordination of theta phases necessary for successful working memory function? To examine this, we shifted the preferred phases of MECII neurons by 180 degrees from the troughs to the peaks of the reference theta oscillation of CA1 LFP (Figure S2A). Figure S2B shows the phase preferences of CA3, CA1 E, CA1 PV, and MECIII E neurons after this change. MECIII PV neurons dramatically reduced spikes at the descending phases of theta oscillation, which shifted the firing of MECIII E neurons to the troughs of theta oscillation. Consequently, the peak activities of MECIII and CA3 were separated by about one half of theta cycle and did not coincidently innervate CA1. The timing deviation impaired the encoding of spatial information into the loop circuit, as indicated by significantly reduced rate differences between L and R subgroups (Figure S2C, p= 2.679×10^−16^, *t* test on two related samples), resulting in a degraded task performance (Figure 2D, p= 2.334 ×10^−8^, *t* test on two related samples). Thus, the specific coordination of the preferred theta phases of MECIII and CA3 neurons is crucial for the working memory operation of the entorhinal-hippocampal circuit.

### The role of disinhibition in regulating the activity of the CA1-MECV-MECIII loop circuit

We show that the ACh-mediated disinhibitory mechanisms regulate cross-area communications within the entorhinal-hippocampal circuit during different task periods. We first analyzed how activity of CA1 E neurons is regulated by [ACh]. We consider the default stage (i.e., 0 to 1 sec in Figure 2B) in which [ACh] is low (Figure 3A). In this stage, output from VIP neurons is weakened and, consequently, PV and OLM neurons are strongly activated around the peaks and the troughs of theta oscillation, respectively (Figure 3B). Accordingly, CA1 E neurons rarely fire around the peaks and troughs (but they can generate a small number of spikes driven by external noise after the troughs of theta oscillation at which inputs from both PV and OLM are weakened: see Figure S1). In contrast, during the epochs of high [ACh] (Figure 3D, the sample-L period: the test-C period also corresponds to the high [ACh] epoch, but will be discussed later), CA1 E neurons show strong activation immediately after the peaks of theta oscillation because PV neurons are suppressed around the peak (Figure 3E). OLM neurons are also suppressed, but their inhibitory effect on E-neuron firing around the peak is relatively weak since the preferred phase of OLM neurons is the trough of theta (spikes around the peak in Figure 3E are less than those in Figure 3B). Thus, [ACh] regulates the activity of CA1 E neurons. Also, the cholinergic modulation advances the preferred phase of CA1 E neurons from the troughs to the peaks of theta oscillation. Later, we will examine the model’s prediction in experimental data.

**Figure S2 (related to Figure 2).**
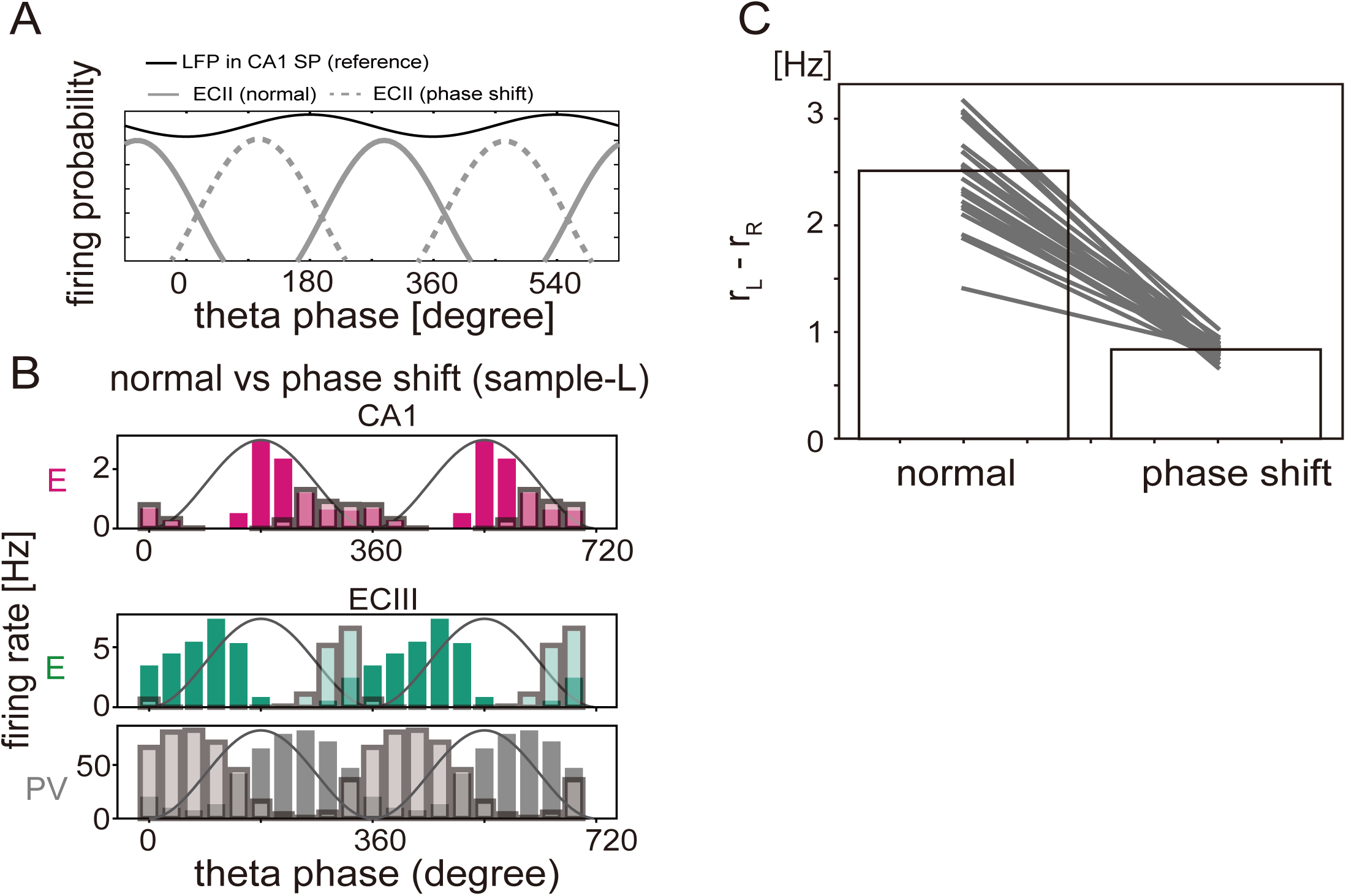
(A) Theta-modulated firing probabilities of ECII neurons for the normal and preferred phase shift conditions are shown. (B) Modulations of spike counts by theta oscillation are shown for CA1 E, MECIII E and MECIII PV neurons during the sample-L period. Neuronal activities in the normal and preferred phase shift conditions of theta oscillation are shown with dark and light colors, respectively. (C) Differences in the firing rate between the L and R subgroups of CA1 E neurons were calculated during the sample-L period and compared between the normal and the phase shift conditions. Simulation results for the same initial conditions are connected with lines.

**Figure 3.**
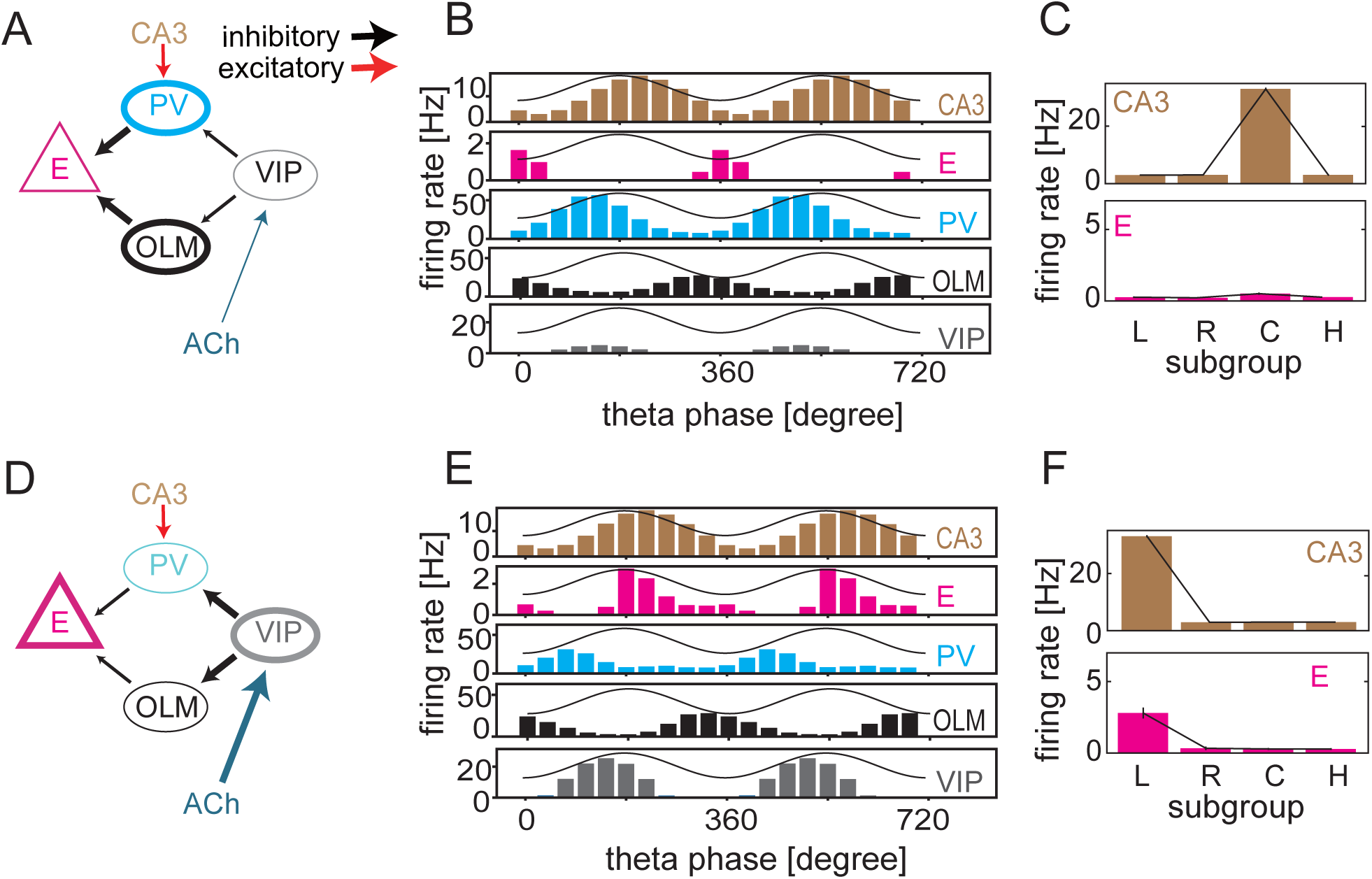
Gating of CA3-to-CA1 signaling by disinhibition mechanism. (A) The operation mode of disinhibitory circuit is schematically illustrated in low [ACh] states. (B) Theta phase preferences of spikes during the sample-C period are shown for CA1 neurons including all subgroups. Only for VIP neurons, the firing rate normalized by [ACh], which corresponds to output from VIP neurons to other inhibitory neurons (see the Materials and methods), is plotted. Solid curves show the reference theta oscillation. (C) Average firing rates of the L, R, C and H subgroups in CA3 and CA1 are shown during the sample-C period. Error bars indicate s.d. (D) The operation mode of disinhibitory circuit in high [ACh] states. (E, F) Similar to (B) and (C) during the sample-L period.

Next, we asked if Information on the current position of the mouse can be selectively transferred from CA3 to CA1 by [ACh]. Although, throughout the task, CA1 E neurons constantly receive position information from CA3, a certain mechanism is required to transfer CA3 position information to CA1 neurons selectively during encoding epoch (in the sample-L period, see the CA1 L subgroups in Figure 2B). We found that ACh-induced disinhibition provides this mechanism: at the sample-C period [ACh] is low and the sensitivity of CA1 E neurons to CA3 input remains low, consequently the position information is not transferred to CA1 (Figure 3C); at the sample-L period [ACh] is increased and accordingly the sensitivity is also enhanced, resulting in an information transfer (Figure 3F).

The disinhibition mechanism further explains why the blockade of MECIII-to-CA1 connections impaired task performance in experiment (Yamamoto et al., 2014). When the model mouse is sampling left or right arm, the disinhibition mechanism enables the activation of the corresponding subgroups in the CA1-ECV-ECIII loop: highly activated CA1 neurons activate MECV E neurons, which in turn activate MECIII E neurons. MECIII-to-CA1 projections further activate CA1 E neurons, thus completing a positive feedback loop within the activated subgroups. However, the blockade of MECIII-to-CA1 connections reduces neural activity in CA1 and hence in MECIII, disabling the storage of the current position.

To confirm the crucial roles of disinhibition (i.e., VIP-PV-E and VIP-OLM-E connections) in the working memory task, we studied two cases. In the first case, PV neurons were inactivated in CA1 during the entire trial period without changing the other conditions (Figure S3A). Due to the lack of inhibition from PV neurons, CA1 E neurons were strongly activated at any ACh concentration. In the encoding epoch, E neurons exhibited higher activity in the L subgroup than in the R subgroup in both CA1 (Figure S3B) and ECIII (Figure S3C, 1 to 3 sec). However, after this epoch E neurons immediately lost selectivity to spatial information because they were too strongly activated in both subgroups. During some intervals firing rate was higher in the L subgroup than in the R subgroup, but it was opposite during other intervals (Figure S3C, after 3 sec). Thus, the spatial information recalled randomly varied from trial to trial, and working memory performance was unreliable (Figure S3D).

In the second case, the cholinergic modulation of PV neurons (but not that of OLM neurons) was terminated during the entire trial period. In this case, PV neurons were not inactivated even at high [ACh] (Figure S3E) and the encoding of spatial information into CA1 was largely impaired (Figure S3F). Consequently, spatial information could not be stably maintained in MECIII (Figure S3G) and probability of left choice, *P*_*L*_, was significantly decreased (Figure S3H, p=1.542×10^−9^, *t* test on two related samples). Thus, in both cases, spatial information in CA3 was not successfully transferred into the CA1-MECV-MECIII loop circuit.

### Covert activation by calcium dynamics

An unexpected experimental finding was that neural activities in MECIII were strongly suppressed during delay periods (Yamamoto et al., 2014). This observation challenges our hypothesis that either MECV or MECIII, or both, serve for working memory in the spatial decision making task, raising the question about how the entorhinal-hippocampal circuit recalls the encoded spatial information after the delay periods. To explore the underlying circuit mechanisms of memory recall, we implemented calcium-sensitive non-specific cation current (CAN current in Materials and methods) in our cortical neuron models.

**Figure S3 (related to Figure 3).**
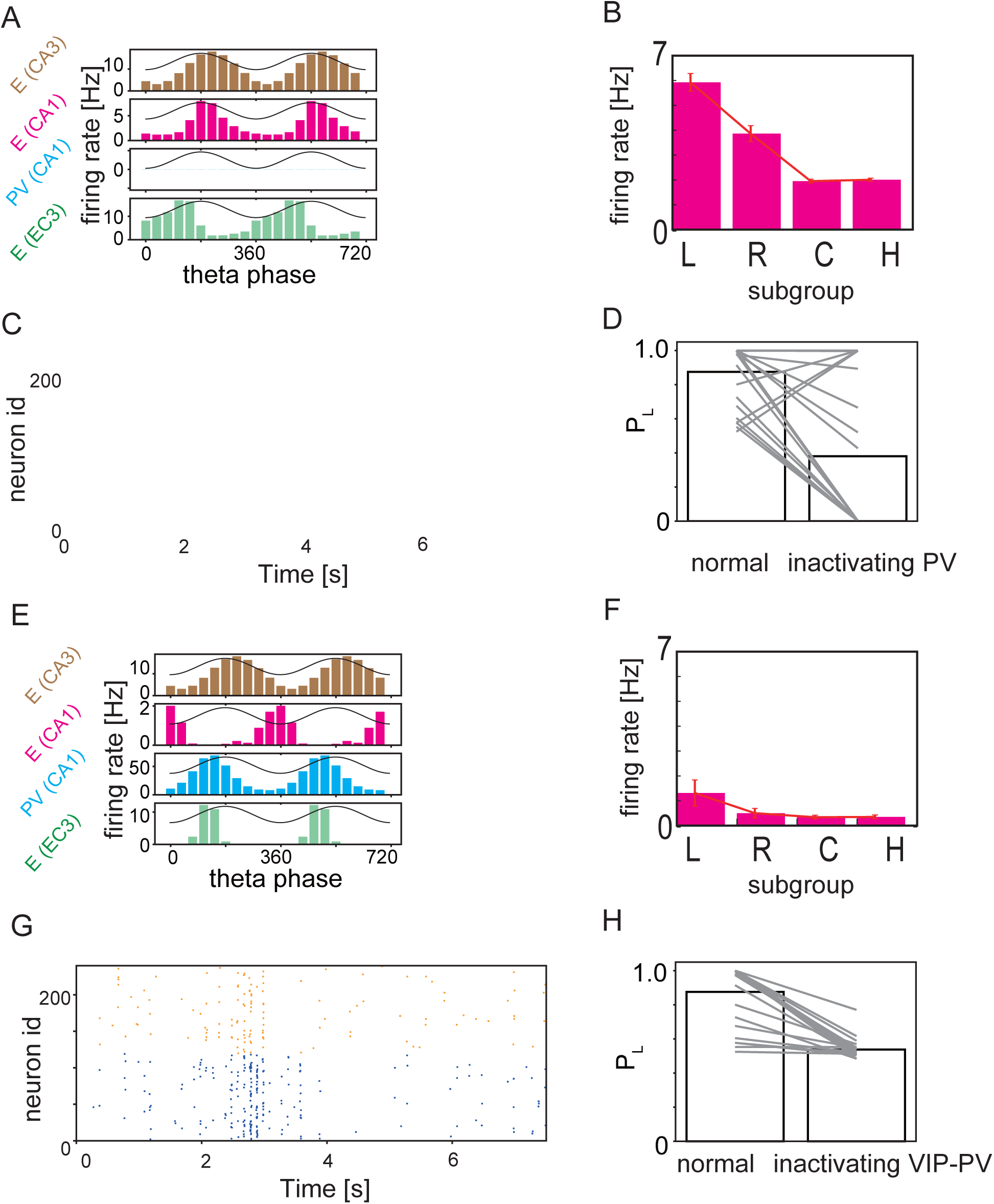
Effect of inactivation of disinhibition mechanism. (A) Firing rates of CA1, CA3 and MECIII neurons during sample-L periods are shown. During the simulations, CA1 PV neurons were inactivated. Solid curves indicate the reference theta oscillation. (B) Average firing rates of CA1 E neurons in different subgroups are plotted during sample-L periods in the same inactivating condition. (C) Raster plots are shown for the L (blue) and R (orange) subgroups of MECIII E neurons. (D) Probability of left choice P_L is plotted in the normal and inactivating conditions. Lines connect two data points obtained from simulations of a normal network and its impaired version with the same initial conditions. (E, F, G, H) same as (A) to (D), but for the network models with disabled VIP-to-CA1 PV connections.

The CAN current was originally proposed to explain persistent activity of single cortical neurons in MECV (Fransen et al., 2002; Fransén et al., 2006), and a similar persistent activity was later shown in the layer V of various cortical areas (Rahman & Berger, 2011). The CAN current is activated in the presence of ACh with the intensity depending on the activation rate of high conductance channels, *r*_H_. The conductance *r*_H_ increases in time when the calcium concentration [Ca^2+^] is beyond a critical value *d*_P_ and decreases when [Ca^2+^] is below another critical value *d*_D_. Because *d*_D_ < *d*_P_, a hysteresis effect or bistability appears for *d*_D_ < [Ca^2+^] < *d*_P_. Thus, once [Ca^2+^] exceeds *d*_P_, the value of *r*_H_ remains high until [Ca^2+^] again decreases below *d*_D_ (Figure 4A). Neurons with high *r*_H_ respond to input more sensitively than those with low *r*_H_ and, thus, memory of previous high activity is stored in *r*_H_ through calcium dynamics.

**Figure 4.**
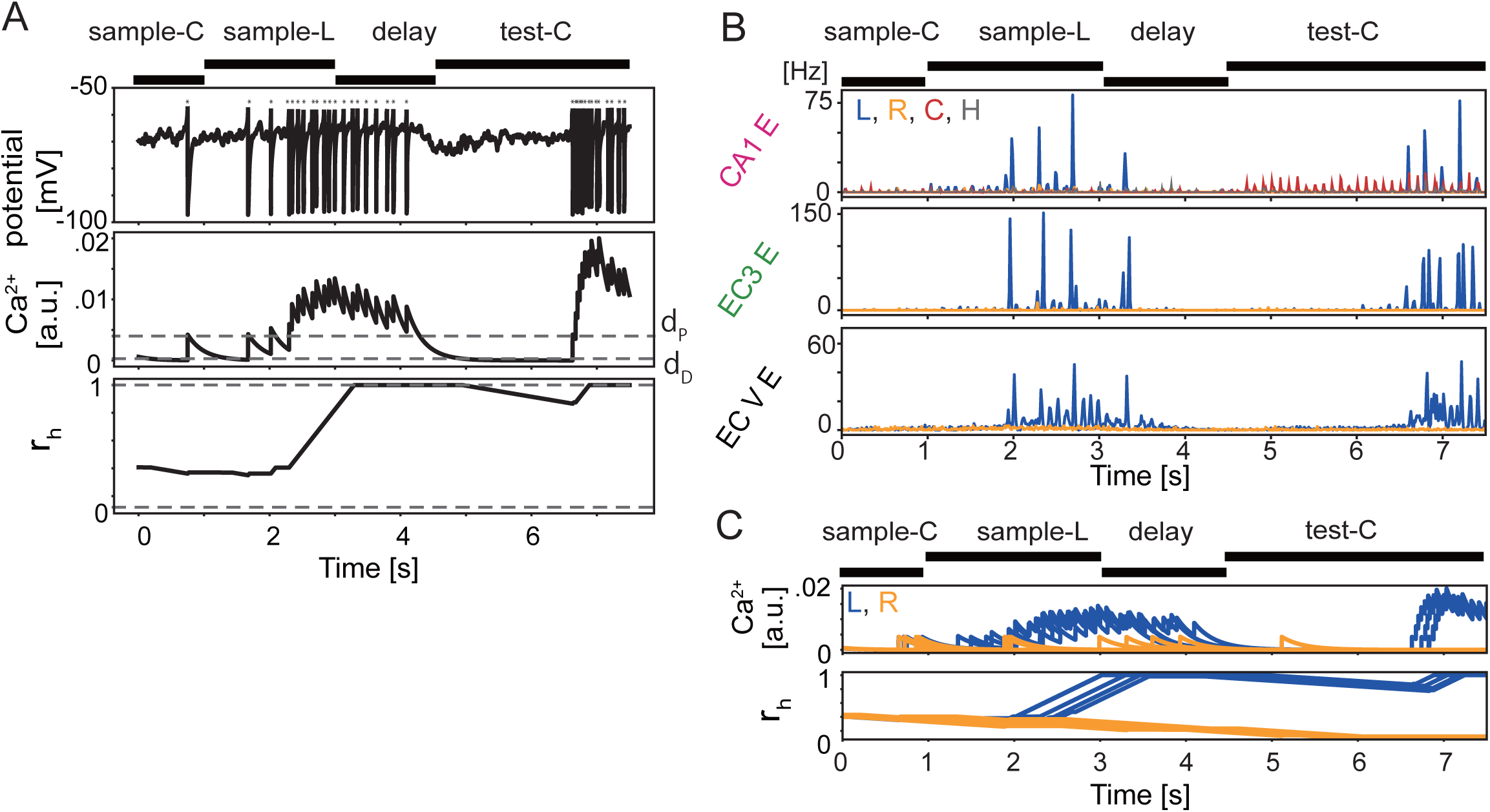
Role of CAN current in memory encoding and maintenance. (A) Single trial evolution of the membrane potential (top), [Ca2+] (middle) and the ratio of high conductance state of CAN channels r_h (bottom) are plotted in an MECV excitatory neuron. Dots above the membrane potential represent spikes. Broken lines denote two threshold values, dP and dD, in the middle panel and the upper and lower critical values of the high conductance ratio in the bottom panel (See Materials and methods). (B) Firing rates are shown for E neurons in CA1, MECIII and MECV. Colors indicate different neuron subgroups. (C) Time evolution of [Ca2+] (top) and rh (bottom) are plotted for randomly-chosen five MECV E neurons belonging to L (blue) or R (orange) subgroup during the same trial as in B.

In the sample-L period, an increased activity of the CA1 subgroup L enhanced spike firing of MECV E neurons in the subgroup L (Figure 4B). During the enhanced firing, [Ca^2+^] was elevated in these neurons by calcium influx through the voltage-dependent calcium channel. This increase of [Ca^2+^] occurred only in the MECV subgroup L but not in the MECV subgroup R (Figure 4C). After the sample-L period, [ACh] was decreased and, consequently, the CAN current was also decreased. Because lowering [ACh] decreased the output of VIP neurons, that of PV neurons was increased and consequently that of CA1 E neurons was suppressed. Thus, the changes in neural activity resulted in a decreased firing rate of MECV E neurons. Nevertheless, the fraction of high conductance state remained high in the subgroup L (but not in the subgroup R) of MECV E neurons (Figure 4C).

The CAN current plays a crucial role in the maintenance of working memory. To explain this, we divided the test-C period into early and late periods: in the early period CA1 neurons were selectively activated in the subgroup C but not in the subgroup L (and subgroup R); in the late period they were strongly activated in the subgroup L (Figure 4B). During the test-C period, [ACh] was again increased, so was the activity of CA1 E neurons through the disinhibition mechanism. Although MECV neurons in both subgroups L and R received synaptic input from the CA1 subgroup C, MECV neurons were selectively activated in the subgroup L because the high conductance rate remained high in these neurons. The activity of the MECV subgroup L neurons gradually increased in the early test-C period, and eventually became sufficiently strong to activate MECIII subgroup L neurons. Accordingly, the test-C period entered the late period and the spatial information stored in MECV could be decoded by CA1 neurons. The onset time of the late period depends on the realization of neural networks and initial conditions. Thus, the covert activation of CAN current enables the retrieval of persistent activity in the MECV subgroup L neurons for the decoding of spatial information in the test-C period.

### Comparison of MECIII neural activity between the model and experiment

We compared the responses of our model with those of the mouse entorhinal-hippocampal circuit. For this comparison, we analyzed E neuron activity in MECIII after dividing each of the sample-L, delay and test-C periods into early and late portions, respectively. As in experiment (Yamamoto et al., 2014), we first quantified the intensity of theta oscillation in these task periods by computing periodicity index (Materials and methods). As shown in Figure 5A, this index was high during the late sample-L, early delay, and late test-C periods, but it was low during delay period (i.e., late delay and early test-C periods). Periodicity index exhibited similar task period-dependence in the model and experiment (c.f. Figure 5C and Figure S5 in (Yamamoto et al., 2014)).

**Figure 5.**
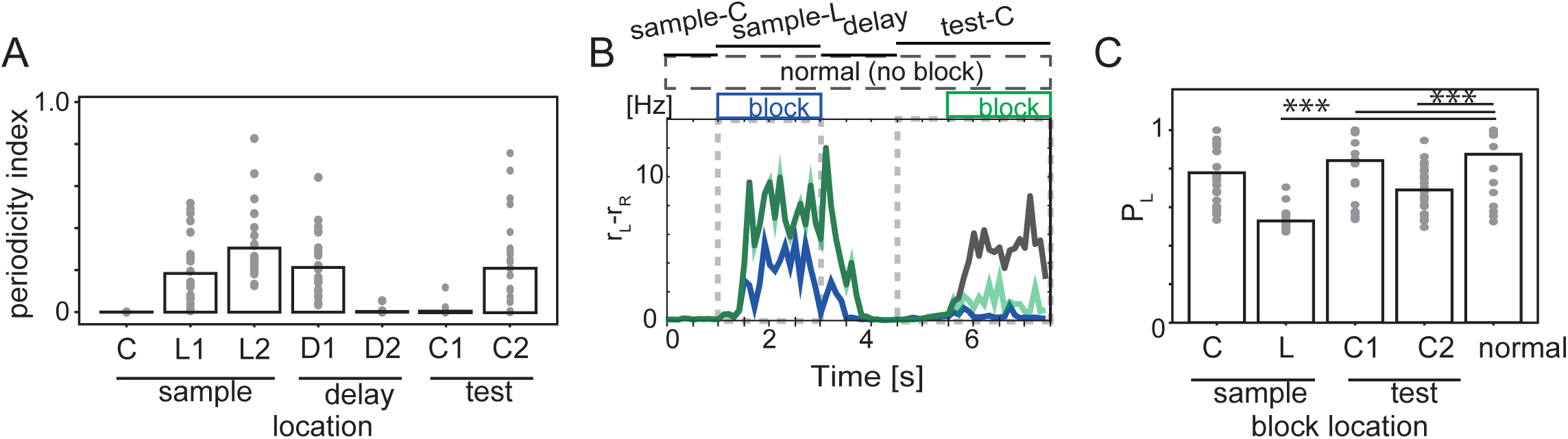
Blockade of MECIII-to-CA1 connections during sample-L and late test-C periods. (A) Periodicity index (Yamamoto et al., 2014) was calculated for the activity of MECIII E neurons (Materials and methods). The labels L, D, C refer to sample-L, delay, test-C periods, respectively, and the numbers 1 and 2 label the early and late portions, respectively, of these periods. (B) Average differences in firing rate between the L and R subgroups of MECIII E neurons were calculated under the blockade of MECIII-to-CA1 connections: normal condition (black); the blockade during sample-L periods (blue); the blockade during late test-C periods (green). In each condition, the differences were averaged over five networks and five initial states. By definition, green and black lines overlap with one another before the blockade. Boxes above the traces indicate the periods of blockade. (C) P_L were calculated for different periods of the blockade.

We next analyzed how the blockade of MECIII-to-CA1 projection affects the behavior of our model in different task periods. In the experiment, this blockade significantly impaired working memory performance of the mouse. When MECIII-to-CA1 projection was blocked during the encoding epoch (sample-L period), MECIII activity and the firing rate difference between L and R subgroups were suppressed during both sample-L and the subsequent late test-C periods (Figure 5B, blue), meaning that task performance was impaired. When the blockade was during the recall epoch (late test-C, 5.5-7.5 s), the inter-subgroup difference was reduced and task performance was also impaired (green line in Figure 5B). In contrast, when the blockade was imposed during sample-C (3-4.5 s) or early test-C period (4.5-6 s), MECIII activity was not greatly affected (Figure S4). Figure 5C summarizes the resultant task performance of the model. In three of the four conditions (blockade in sample-C, early and late test-C), results were well consistent with experimental observations (c.f., Figure 6F in (Yamamoto et al., 2014)). In addition, our model predicts that the blockade in sample-L period significantly impairs working memory performance (p=3.175×10^−10^), suggesting that the CA1-MECV-MECIII loop circuit maintains neural activity in the MECIII and plays a pivotal role in the spatial working memory task. This prediction should be experimentally validated.

**Figure S4 (related to Figure 5).**
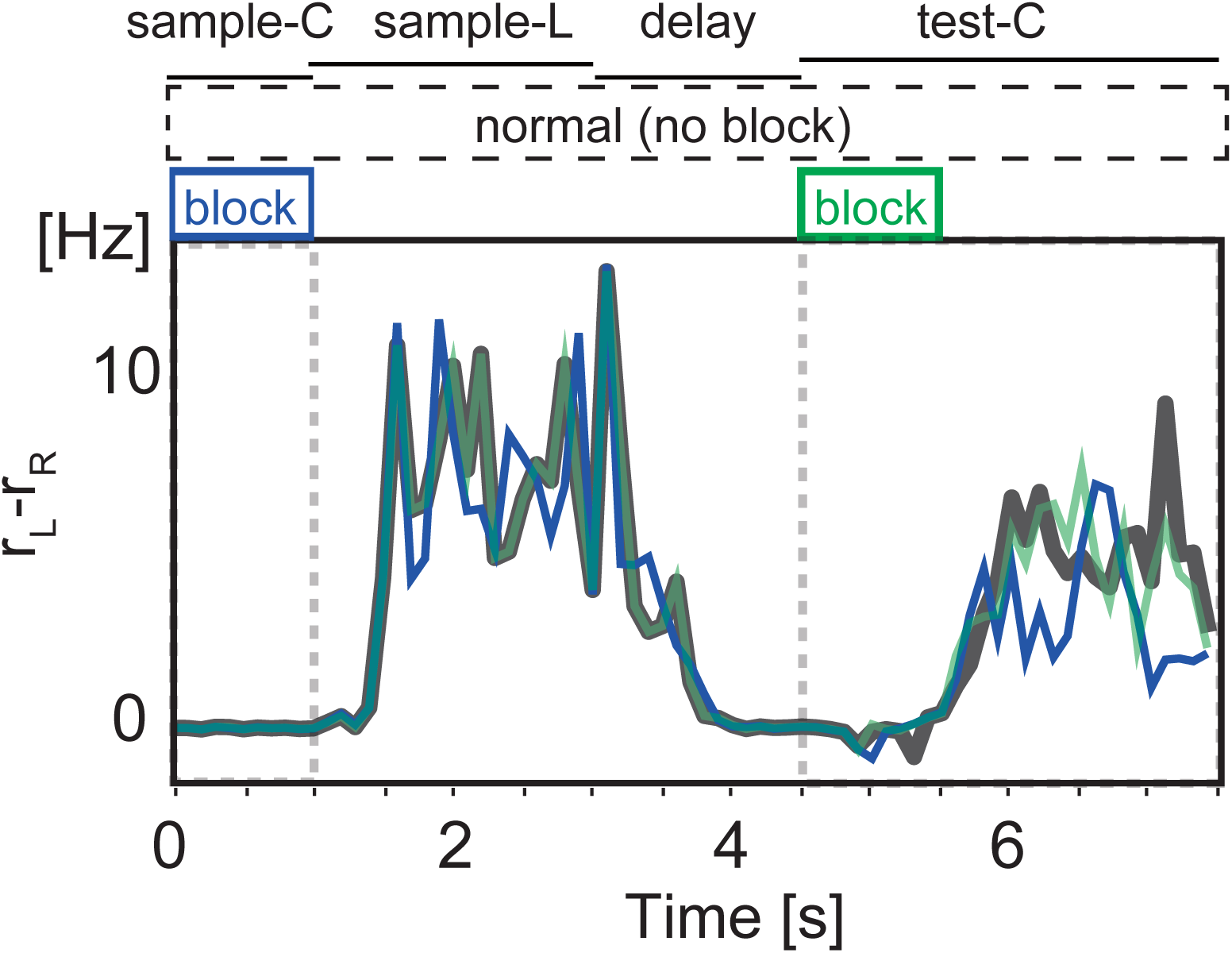
Blockade of MECIII-to-CA1 connections during sample-C and test-C periods. Connections from MECIII to CA1 were blocked during sample-C (blue) and early test-C (green) periods. Differences in firing rate between the L and R subgroups of MECIII E neurons are shown in the same manner as in Figure 5B. Similar evolutions are also shown in the normal condition (black).

**Figure 6.**
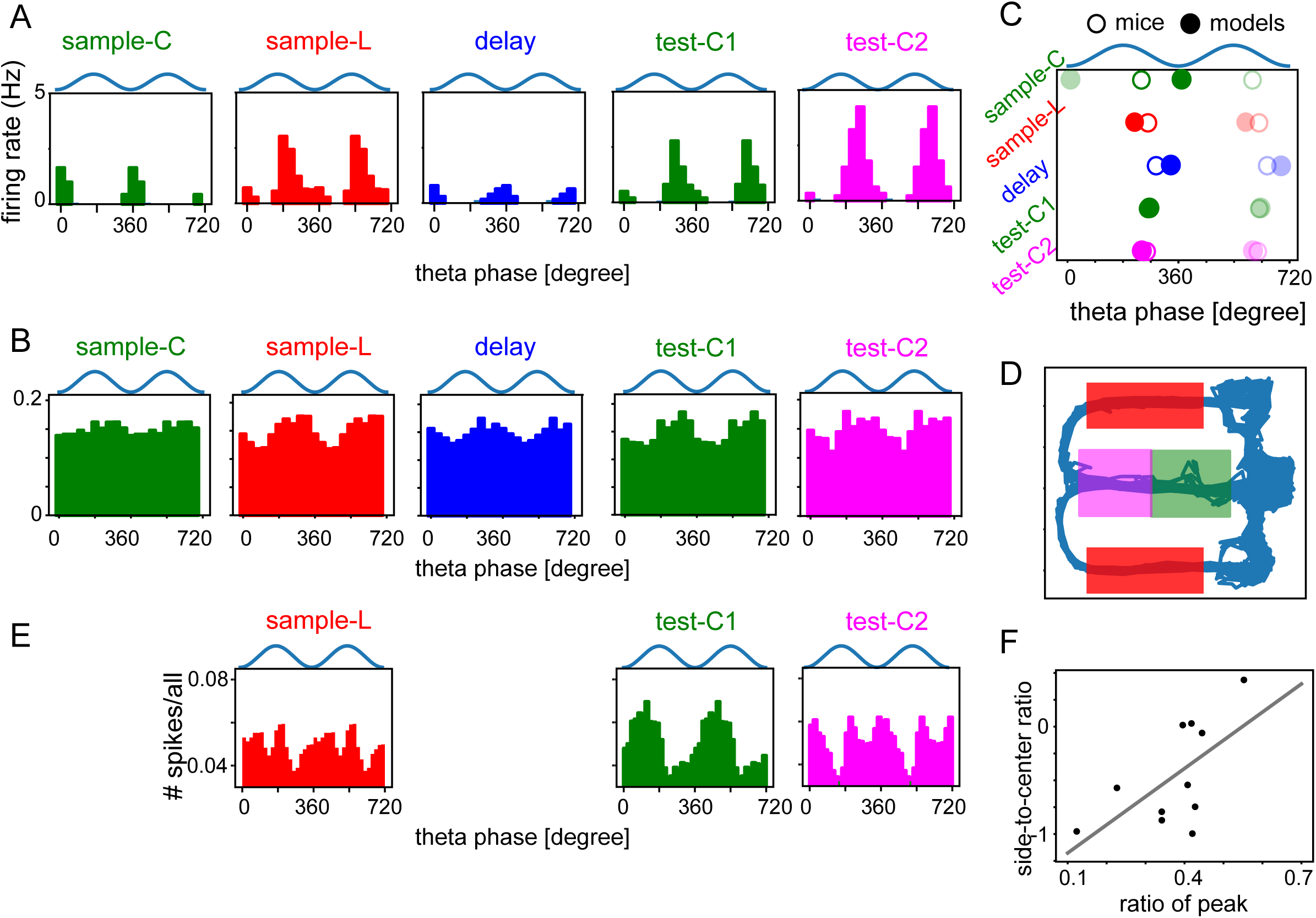
Preferred theta phases in model networks and rodents. (A) Phase preferences are shown for CA1 E neurons in model networks during given task periods (top). (B) Phase preference curves of CA1 neurons were calculated for data obtained in (Yamamoto et al., 2014). (C) Preferred theta phases of the models (filled circles) and mice (empty circles) were averaged over multiple theta cycles during given task periods. The same average phases are shown for two theta cycles (one in darker colors and one in light colors) for the clarity of the plots. (D) Schematic illustration of the figure-eight maze used in (Fernández-Ruiz et al., 2017; Mizuseki et al., 2013). Colored rectangles indicate early-center, late-center, and reward periods from which data were resampled. (E) Phase preference curves of CA1 excitatory neurons were calculated by using the data of (Mizuseki et al., 2013). For comparison, we divided the task periods of center arm and reward arm into early and late epochs. (F) Correlation between the ratios of spikes at the peaks and the side-to-center ratio is plotted.

### Experimental validation of period-dependent preferred theta phases

Our model predicts that CA1 E neurons shift their preferred theta phases from the troughs to the peaks when [ACh] is high (Figures 3B and 3E). In Figure 6A, we present the phase preference of CA1 E neurons during sample-C, sample-L, delay, early test-center (test-C1) and late test-center (test-C2) periods of the DNMP task. Their spikes preferred the troughs of theta oscillation during sample-C periods but, owing to the disinhibitory mechanism, the preferred phase was shifted to the peaks during sample-L periods. When the model mouse returned to the home position (delay period), [ACh] was decreased to reactivate PV neurons in CA1, which reduced the sensitivity of CA1 to inputs from MECIII and CA3 and selectively suppressed spike generation at the peaks of theta oscillation (but not at the troughs). In test-C1 periods, [ACh] was again increased to allow the activation of CAN current (Figure 4C), which shifted the preferred phase to the descending phase of theta oscillation. During test-C2 periods, our model predicts a progressive advance of preferred theta phase in CA1 due to an enhanced synaptic drive by MECIII.

We confirmed these predictions in the data of a DNMP task in T-maze (Yamamoto et al., 2014). Figure 6B shows the distributions of preferred phases of spikes in mouse CA1 during different task periods, i.e., sample-C, sample-L (corresponding to the reward arm), delay, test-C1, and test-C2 periods (see Materials and methods). In mice, spikes were generally not modulated by theta oscillation as strongly as in the model. In particular, oscillatory modulations were weak in sample-C periods. Nonetheless, the preferred theta phases in the various task periods are well consistent between mice and models. The average preferred phase, which was computed as 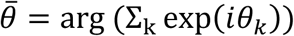 with k being the index of spikes for all neurons and *i* being imaginary unit, is delayed during delay periods compared to other task periods in both mice and models (Figure 6C). In contrast, our model predicts that the preferred phase is progressively advanced during test-C2 (i.e., late test-center) period due to an increased synaptic drive by MECIII (Figure 4B).

Next, we asked whether preferred theta phase behaves similarly in an alternating figure-eight task (Mizuseki, Sirota, Pastalkova, Diba, & Buzsaki, 2013). We were particularly interested in examining the hypothesized role of cholinergic control of working memory function. In the alternating figure-eight task, rats were trained to alternately change the turn direction at a junction point of an eight-shape maze, meaning that the rats had to remember the turn direction of the preceding run. This task is similar to the previous DNMP task, but one difference is that sample and test trials are not clearly separated in the alternating figure-eight task. Nevertheless, we may correlate the behavioral epochs of the two tasks to each other. When rats traverse the center arm, they had to retrieve memory of the preceding choice to prepare for the next choice. Therefore, traveling along the center arm may correspond to test-C period in the DNMP task. Then, we can define three distinct areas on the eight-shape maze (Figure 6D): early center, late center and reward arms, which may correspond to test-C1, test-C2 and sample-L periods, respectively. Below, we follow these rules.

Figure 6E shows the preferred phases of excitatory neurons in the deep layer of CA1. It has been reported that for some unknown reason these neurons only exhibit phase shifts in early trials (Fernández-Ruiz et al., 2017; Mizuseki et al., 2013). Therefore, we only used data of initial ten trials in the following analyses. On the early center arm (test-C1 period), neurons fired more frequently around the troughs of theta oscillation than the peaks. However, on the late center arm (test-C2 period), neurons fired slightly more often at the peaks, generating two peaks per theta cycle in the spike density distribution. On the reward arm (sample-L period), neurons fired most frequently at the peaks, which is consistent with the previous study(Fernández-Ruiz et al., 2017). These results seem to be consistent with the model’s prediction that the preferred firing phase of CA1 neurons changes from the troughs to the peaks of theta oscillation during the epochs of high [ACh]: in the alternating figure-eight task, high level of attention, or high [ACh], is likely to be required on the reward arm (for encoding reward information) and the late center arm (for recalling the previous choice).

Finally, we explored single-cell-level behavior in (Mizuseki et al., 2013) to provide further support for our prediction: cells spiking on the side arms (cells in L or R subgroup in the model) are likely to spike on the late center arm at the peaks of theta oscillation. Therefore, we computed two ratios for 11 cells which showed high activation on the late center arm (See Materials and methods for details): the number ratio of spikes generated around the peaks (90° to 270°) to spikes generated on the late center arm; the number ratio of spikes generated on the side arms to spikes generated on the late center arm (side-to-center ratio). The values of the two ratios are significantly correlated (Figure 6F: r=0.62 and p=0.042, *t*-test), suggesting that cells strongly activated on the side arms are also highly likely to fire around the theta peaks. This result strongly supports the prediction of our model.

### Task performance depends on acetylcholine concentrations in the model

In our model, two ACh-regulated mechanisms, that is, disinhibitory circuit in CA1 and CAN current in MECV excitatory cells, play crucial roles in encoding, maintenance and recall epochs of working memory tasks. Therefore, we examined whether these core mechanisms work robustly when [ACh] is changed. We changed the default levels of [ACh] with other parameter values unchanged. Lowering the default [ACh], which weakens disinhibition, during the encoding epoch (sample-L) disenabled the CA1 L and R subgroups to exhibit large enough activity differences to encode spatial information in the CA1-MECV-MECIII loop circuit (Figure 7A, blue). Task performance was severely impaired, contrasting to intact performance at higher default [ACh] levels (Figure 7B, left). Lower default [ACh] levels during delay period had almost no effects on task performance (Figure 7B, center). Finally, lower default [ACh] levels during the recall epoch (test-C2) eliminated the reactivation of the subgroup L (Figure 7A, green) and task performance was severely impaired (Figure 7B, right), but performance remained intact at higher [ACh] levels. These results show that default [ACh] levels should be sufficiently high during the encoding and recall epochs, but fine tuning of [ACh] is unnecessary.

**Figure 7.**
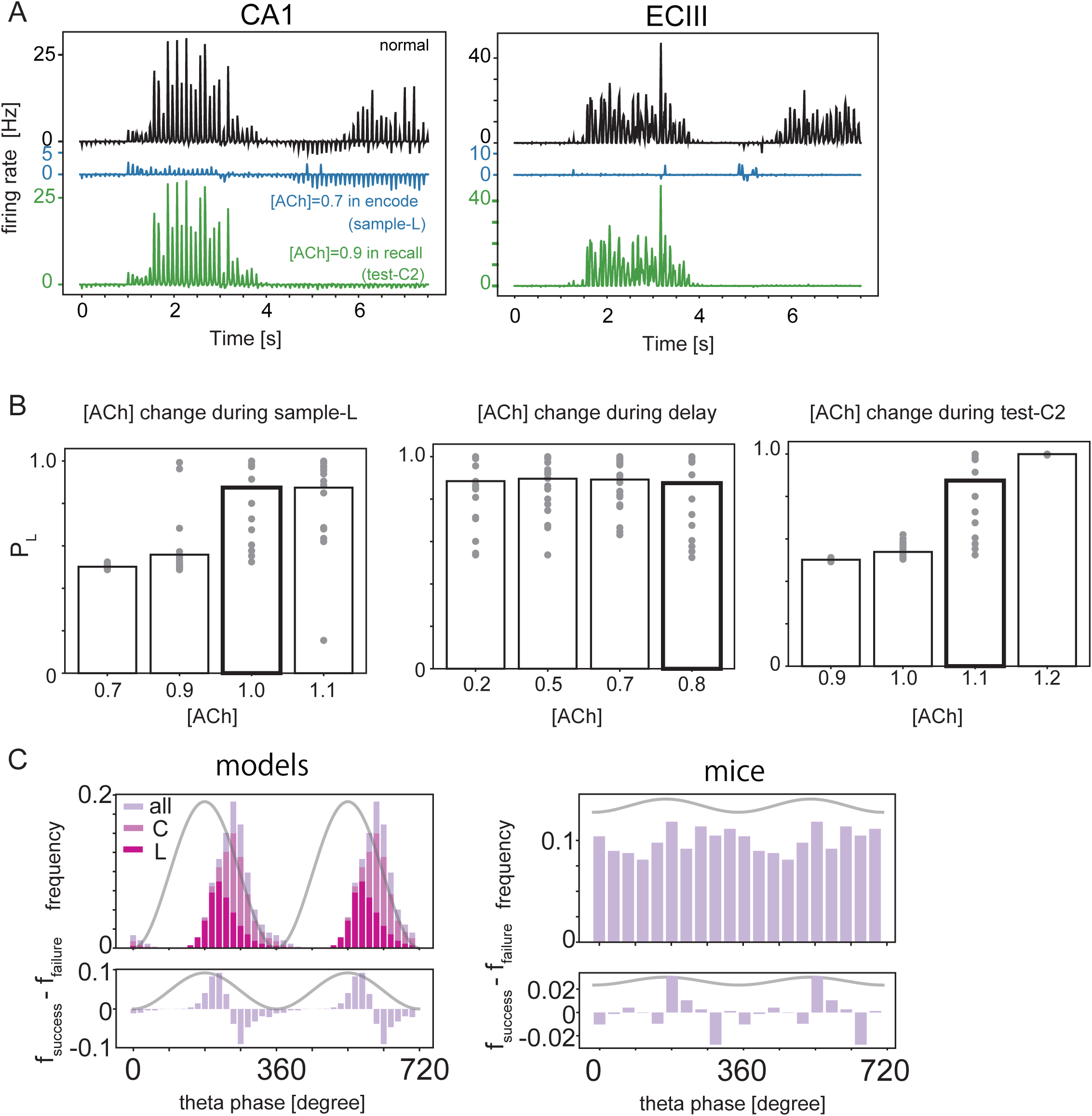
Cholinergic modulations in the network model. (A) In CA1 (left) and MECIII (right), differences in firing rate between the subgroups L and R were calculated at the normal (black) and reduced levels of [ACh] during encoding (sample-C, blue) and recall (test-C2, green) epochs. (B) Probability of left choice averaged over different networks and initial conditions are shown at different levels of [ACh] during sample-L (left), delay (center) and test-C2 (right) periods. Thick lines indicate results for the default [ACh] levels. (C) Frequency of spikes in CA1 excitatory neurons during test-C2 periods (top) are shown in the model (left) and mice (right). In the model, [ACh] was set to the normal level and spike counts are shown separately for the subgroups L and C. Only the total spike count is shown for mice. Bottom panels show the differences in firing rate between successful and failure trials.

Next, we explored whether theta-phase-locked firing has anything to do with the success of the working memory task. To this end, we compared the theta preference of CA1 neurons during the test-C2 period between successful and failure trials in both models and mice (Figure 7C). [ACh] was at the default levels. In both models and mice, neurons preferentially fired around the peaks of theta oscillation in success trails, but firing phases were somewhat delayed in failure trials.

## Discussion

In this study, we developed a model comprising MECV, MECIII and CA1 to explore how these local circuits process and communicate information with each other regulated by ACh during DNMP tasks. In our model, changes in the ACh concentration control cortical disinhibitory systems and calcium-dependent cationic current to perform different cognitive functions in DNMP tasks. With the ACh modulations, our model successfully replicates the various features of neural activity observed in MECIII and CA1 (Yamamoto et al., 2014). In particular, the model predicts that CA1 neurons change their preferred theta phases depending on cognitive demands, which was also supported by experimental data (Fernández-Ruiz et al., 2017; Mizuseki et al., 2013; Yamamoto et al., 2014).

### Relevance of MEC-CA1 loop to spatial working memory tasks

The hippocampal area CA1 is a central locus for spatial information processing, and stores the concurrent position of the rodent (O’Keefe & Dostrovsky, 1971) as well as retrospective and prospective representations of position information (Dragoi & Buzsáki, 2006; Ferbinteanu & Shapiro, 2003; Foster & Wilson, 2006; Gupta, van der Meer, Touretzky, & Redish, 2010; Pastalkova, Itskov, Amarasingham, & Buzsaki, 2008; Zheng, Bieri, Hsiao, & Colgin, 2016). Focusing on the CA1-MECV-MECIII loop circuit involved in spatial working memory (van Strien, Cappaert, & Witter, 2009; Witter et al., 2000), we demonstrated how the spatial information of the selected arm is encoded in CA1 during a sample trial (Figure 3), maintained in MECV during delay period, and transferred to MECIII and reloaded on CA1 for decision making (Figure 4). Several studies found that connections from MECIII to CA1 are crucial for the success of spatial working memory tasks (Suh et al., 2011; Yamamoto et al., 2014). Our model demonstrated this role of MECIII-to-CA1 connections in spatial working memory, predicting their crucial contribution to memory encoding by the entorhinal-hippocampal loop circuit. The predicted operation of the loop circuit should be tested experimentally.

Sequence of spikes with theta phase precession, that is, “theta sequence” (Ferbinteanu & Shapiro, 2003; S. J. Middleton & McHugh, 2016; Pfeiffer & Foster, 2013; Schlesiger et al., 2015; Zheng et al., 2016), has been observed in the hippocampus and the MEC during cognitive tasks requiring episodic memory. The blockade of MECIII-to-CA1 connections causes modulation of theta activity in MECIII (Suh et al., 2011; Yamamoto et al., 2014).

Similarly, inactivation of the MEC disrupts the temporal organization of spikes and impairs information maintenance (Robinson et al., 2017). These results imply that the temporal coordination of theta-phase-locked neuronal firing along the CA1-MECV-MECIII loop circuit is crucial for the success of spatial working memory tasks. We showed an example case in which the disruption of this temporal coordination in MECIII led to a significant increase of failure trials (Figure S2A, B).

Our model assumes that [ACh] changes in time during a working memory task and predicts that these changes shift the preferred theta phase in CA1 E neurons: higher [ACh] advances the preferred phase towards the peak of theta oscillation (Figure 6). Specifically, the model predicts that phase advances occur during the encoding epoch in sample trials (i.e., on the reward arm) and during the recall epoch in test trials (i.e., on the center arm). We validated this prediction with the results of two experiments (Fernández-Ruiz et al., 2017; Mizuseki et al., 2013; Yamamoto et al., 2014). Phase advances in the encoding epoch have been known previously and can be accounted for by temporal separation between CA3 input and MECIII input to CA1 (Colgin et al., 2009; Cutsuridis, Cobb, & Graham, 2010; Hasselmo, Bodelón, & Wyble, 2002; Lasztóczi & Klausberger, 2016; Milstein et al., 2015). Input from CA3 preferentially arrives at CA1 on the descending phase of theta oscillation of LFP (Klausberger & Somogyi, 2008) whereas input from MECIII arrives at the peaks of theta oscillation (Fernández-Ruiz et al., 2017; Hasselmo et al., 2002; Mizuseki et al., 2009). Because MECIII input, which presumably carries sensory information, seems to dominate CA3 input during encoding (Hasselmo2002), CA1 neurons likely fire at the peaks rather than the troughs of theta oscillation (Fernández-Ruiz et al., 2017).

Phase advances on the later central arm (i.e., in the recall epoch) represent a novel finding of this study. CA1 neurons were previously shown to fire at the troughs of theta oscillation during memory recall (Fernández-Ruiz et al., 2017), and this was consistent with the view that CA3 input dominates MECIII input during this epoch. Challenging the conventional view, our model predicts that CA1 neurons retrieve spatial memory from the MECIII, hence firing at the peaks of theta oscillation. Possibly consistent with this prediction, if we divide the center arm into early and late portions in (Mizuseki et al., 2009), preferred theta phases show a second (and advanced) peak in the late portion of recall epoch (Figure 6D). This peak was absent in the previous analysis (Fernández-Ruiz et al., 2017) because the center arm was treated as a single entity. The consistency between the model and experiment requires further clarification.

### Reflection of cognitive demands on the MEC-CA1 loop circuit through ACh

ACh is involved with several cognitive functions such as sensory discrimination (Hangya et al., 2015; Pinto et al., 2013), associative learning (Sabec et al., 2018), and spatial (Croxson et al., 2011; Okada et al., 2015) and non-spatial working memory (Furey et al., 2000; Hasselmo & Stern, 2006; McGaughy et al., 2005). In correlation with cognitive states, ACh concentration changes to modulate activity of the specific types of neurons (Muñoz, Tremblay, Levenstein, & Rudy, 2017; Womelsdorf et al., 2014) through muscarinic and nicotinic receptors (Parikh et al., 2007; Teles-Grilo Ruivo et al., 2017; H. Zhang et al., 2010). In associative learning, ACh was suggested to facilitate MECIII-to-CA1 input during the encoding epoch and CA3-to-CA1 input during the decoding epoch (Hasselmo, 2006). We propose a novel role and mechanism of ACh for the functions of the entorhinal-hippocampal loop circuit according to the cognitive demands arising during a spatial working memory task, namely, memory encoding, maintenance and recall. Consistent with the proposal of our model (Figure 2B), it has recently been shown in a DNMP task (Teles-Grilo Ruivo et al., 2017) that [ACh] is significantly higher on the reward arm than in other positions in both sample and test trials, that on the center arm [ACh] tends to be larger in test trials than in sample trials, and that [ACh] is low in delay periods (c.f., Figure 7B).

We assumed that a change in cognitive demands is reflected in a phasic change in [ACh] on the timescale of seconds. However, [ACh] is thought to change in a diffusive and tonic manner on much slower timescales of minutes or hours. Importantly, recent studies have revealed that [ACh] undergoes phasic changes at sub-second and second timescales (Parikh et al., 2007; Teles-Grilo Ruivo et al., 2017; H. Zhang et al., 2010), and such a phasic change in [ACh] is associated with reward or aversive signals (Hangya et al., 2015). Further, task performance is correlated with slower tonic increases in [ACh] during the task period (Parikh et al., 2007), but uncorrelated with phasic changes in [ACh] in the reward arm (Teles-Grilo Ruivo et al., 2017). In our model, task performance saturates above a certain level of [ACh] in the reward arm (Left panel in Figure 7B). Our results suggest that the tonic level of [Ach] expresses an overall bias during each trial and a phasic increase in [Ach] gives a more elaborate modulation reflecting a specific cognitive demand.

### Dynamic processing across multiple areas

Coherence in neural activity between different cortical areas varies with the cognitive state of the brain (Benchenane et al., 2010; Fries, 2015). Furthermore, disruption of a cortico-cortical interaction at different behavioral states can impair task performance differently (Spellman et al., 2015; Yamamoto et al., 2014). These results imply that information flows between cortical areas are differentially routed according to the demand of the on-going cognitive process through the dynamical regulation of corticocortical coherence. Theoretical studies have proposed several mechanisms of information routing based on a balance control between excitatory and inhibitory synaptic inputs (Vogels & Abbott, 2009), disinhibitory circuits (Yang et al., 2016), spontaneous bursts (Palmigiano et al., 2017), and band-pass filtering by a feed-forward inhibitory circuit (Akam & Kullmann, 2010). While these studies focused on the circuit mechanisms of information routing, we addressed how such mechanisms are integrated to perform a spatial working memory task through different cognitive demands (Benchenane et al., 2010; Spellman et al., 2015; Yamamoto et al., 2014). We demonstrated that cholinergic inputs coordinate the encoding and recall functions by modulating the cortical disinhibitory circuit and Ca^2+^-dependent cationic channels in excitatory cells.

Accumulating evidence suggests that disinhibitory circuits play a crucial role in various cognitive tasks such as fear conditioning (Letzkus et al., 2011; Pi et al., 2013) and sensory discrimination (Hangya et al., 2015; Pinto et al., 2013). The dominant interneuron types of the disinhibitory circuits are VIP, SOM and PV inhibitory neurons (Donato, Rompani, & Caroni, 2013; Francavilla et al., 2018; Kamigaki & Dan, 2017; S. Zhang et al., 2014). Among these neurons, VIP neurons express muscarinic receptors and are depolarized by cholinergic input (Bell, Bell, & McQuiston, 2014) and are thought to project more strongly to SOM neurons than to PV neurons in cortical areas (Kamigaki & Dan, 2017; S. Zhang et al., 2014). However, some studies suggest stronger cholinergic modulations of PV neurons in the hippocampus (Donato et al., 2013; Francavilla et al., 2018). As explained below, the cholinergic modulation of OLM neurons also does not strongly influence the firing of CA1 E neurons. Therefore, in this study we mainly analyzed the effect of PV neurons on neural circuit functions. In line with our model’s prediction (Figure S3), optogenetic inhibition of PV neurons impairs performance in spatial working memory (A. J. Murray et al., 2011).

PV neurons which preferentially fire in the descending phase of theta oscillation (Klausberger & Somogyi, 2008) to weaken the effect of CA3-to-CA1 input. In contrast, SOM (OLM in CA1) neurons preferably spike at the troughs of theta oscillation (Klausberger & Somogyi, 2008; Royer et al., 2012) much earlier than the CA3 input. In our model, excitatory MECIII input innervates CA1 preferentially at the theta peaks but rarely at the theta troughs. Therefore, the ACh-induced suppression of OLM neurons does not also enhance the effect of MECIII input on excitatory neuron firing in CA1, making OLM neurons less effective than PV neurons in modulating the CA1 activity. In rats, however, the actual spikes delivered by the MECIII are distributed broadly over a theta cycle (Mizuseki et al., 2009), implying that the suppression of PV and SOM neurons can induce a complex modulatory effect in CA1 pyramidal neurons. Our model predicts that the inactivation of PV or VIP neurons (in this case both PV and SOM are released from the inhibition by VIP neurons) impairs task performance in different ways (Figure S3). This prediction needs to be confirmed by experiments.

### Covert memory state for maintenance of information

Recent studies showed that dynamically evolving neuronal activity can maintain information during a delay period in working memory tasks (J. D. Murray et al., 2017; Stokes, 2015; Wolff, Jochim, Akyürek, & Stokes, 2017). In the DNMP task we studied (Yamamoto et al., 2014), theta phase-locked firing of MECIII neurons was correlated with the success of the task. Nevertheless, this neuronal activity temporarily vanished during a delay period, implying that a non-spiking activity maintains information on the previous choice. We hypothesized that the conductance of a specific ionic channel, i.e., calcium-dependent cationic current, remains in an elevated state to preserve spatial information during delay period. This elevated state is not accompanied by neuronal firing, hence is consistent with experimental observations. This mechanism was originally proposed to account for persistent activity of isolated single neurons (Fransen et al., 2002; Fransén et al., 2006) and suggested to be engaged in temporal association memory (Kitamura et al., 2014). Our model demonstrates that the same mechanism can generate a covert memory state necessary in spatial working memory. This and other mechanisms of covert memory state, for instance, short-term synaptic plasticity (Mongillo, Barak, & Tsodyks, 2008), are not mutually exclusive. However, our mechanism has an important advantage that working memory maintenance is turned on and off by cholinergic modulation depending on the cognitive demand. Thus, our results suggest that neuromodulators are crucial for the flexible control of memory processing by the brain.

### Limitation of the model

First, while our model indicates that a success in spatial working memory tasks requires the adequate preferred theta phases of MEC and hippocampal neurons, experimental results suggest an active role of high-gamma oscillation (60-120 Hz) in working memory tasks (Yamamoto et al., 2014). In our model, theta-phase-locked neuronal firing is sufficient for successful information transfers along the MECIII-CA1-MECV loop circuit and gamma oscillation was not modeled. We speculate that gamma oscillation may significantly facilitate the decoding of information stored in the entorhinal-hippocampal circuit from its outside. The role of gamma oscillation in spatial working memory needs to be further explored.

Secondly, we did not model any mechanism to translate the decoded information into a correct choice behavior under a given rule of decision making (e.g., the alternative of left or right turn). The mPFC is projected to by CA1 and projects back to it via reuniens (Dolleman-Van Der Weel & Witter, 1996; Ito, Zhang, Witter, Moser, & Moser, 2015), and is engaged in spatial working memory (Bolkan et al., 2017; Jones & Wilson, 2005; Spellman et al., 2015). Furthermore, some mPFC neurons exhibit rule-related activities. However, the delay-period activity of mPFC neurons is specific to neither previous nor present location in a DNMP task (Bolkan2017) and the rule-related activities are not location-specific (Durstewitz et al., 2010; Guise & Shapiro, 2017; Preston & Eichenbaum, 2013). Where and how decision rules are processed and how spatial working memory and rule-related activities are integrated are open to future studies.

## Materials and methods

### Neuron models

Our model has two classes of neurons: i) Poisson neurons; ii) Hodgkin-Huxley type (HH) neurons.

#### i) Poisson neurons

There are four types of theta-oscillating neurons (excitatory neurons in CA3 and MECII, VIP neurons in CA1 and GABAergic neurons in MS) and noise neurons. Other types of neurons, that is, excitatory (E) neurons, fast spiking (PV) neurons and OLM neurons in CA1, E and PV neurons in MECIII and E and PV neurons in MECV, are modeled as HH neurons. 40 excitatory and 40 inhibitory noise neurons project to all HH neurons. Firing rates of noise neurons are different depending on the cortical areas modeled: 35 [Hz] in CA1 and in MECIII and 30 [Hz] in MECV.

For theta-oscillating Poisson neurons, we described the theta-oscillating (10Hz) probability of spiking per unit time *P*_*x*_(*t*_1_ < *t* < *t*_1_ + Δ*t*) (*x*=E in CA3 and MECII, VIP in CA1, GABA in MS) as

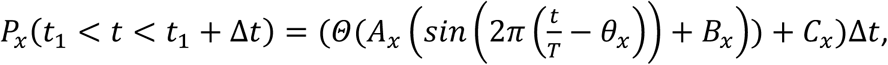

where *Θ(s)* = *s* when *s*>0 or otherwise 0. *T* = 100 ms and *Δ*t = 0.02 ms is the step size of our numerical simulation. *A*_*x*_ and *B*_*x*_ are the amplitude and preferred phase of oscillating firing rate, respectively, and *θ*_*x*_ is the preferred phase of theta oscillation for the neuron type *x*, whereas *C*_*x*_ is the amplitude of background noise for inactive subgroups outside of their place fields. We set the preferred phases based on previous experimental studies (Borhegyi, 2004; Klausberger & Somogyi, 2008; Mizuseki et al., 2009) as follows:

##### *x* = E in CA3

CA3 neurons in the model are divided into four groups according to their spatial preferences (Figure 2 and Circuit structure in Materials and methods). Their firing patterns are changed depending on the present location of the model rat. When the model rat is in Center (0 < *t* < 1000 ms, sample-C and 4500 ≤ *t* < 7500 ms, test-C), sample-Left (1000 ≤ *t* < 3000 ms) and Home (3000 ≤ *t* < 4500ms) positions, Center, Left, and Home subgroups are activated, respectively.

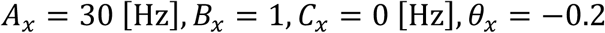

for neurons in an active subgroup, and

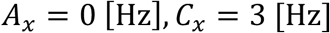

for neurons in inactive subgroups.

##### *x* = ECII

During the entire trial period, we set the parameters as

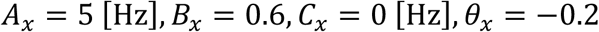

##### *x* = GABAergic in MS

Three groups exist in the model, those projecting to PV in CA1, to OLM in CA1 and to PV in MECV. The preferred theta phase of MECV PV-projecting GABAergic neurons is the same as that of CA1 OLM-projecting GABAergic neurons,

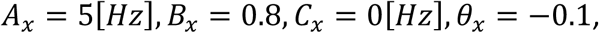

whereas the preferred phases of CA1 PV-projecting neurons are different from the above ones (Borhegyi, 2004) and given by

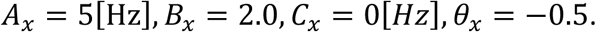

##### *x*=VIP

During the entire trial period, we set the parameters as

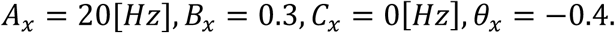

Finally, we set a reference theta oscillation in CA1 SP as

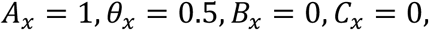

which is a virtual oscillatory component used only for determining the relative oscillation phases of other brain regions to theta oscillation in CA1 SP, but not for numerically simulating the network model.

#### ii) HH neurons

In our model, there are seven types of neurons; E, PV and OLM in CA1, and E and PV in MECV and MECIII. PV neurons are modeled identically in all areas (Wang & Buzsáki, 1996). In the following equations, *I*_*i*_ is synaptic input from other neurons as described in Circuit structure in Materials and methods. Dynamics of each type of neurons is described below.

##### a) Excitatory neurons in CA1

We modeled excitatory neurons in CA1 based on (Wulff et al., 2009). We additionally included an afterhyperpolarization (AHP) and h currents in the model for generating a weak subthreshold oscillation of the membrane potentials through interplay between AHP (S. Middleton et al., 2008) and h current (Rotstein et al., 2006).

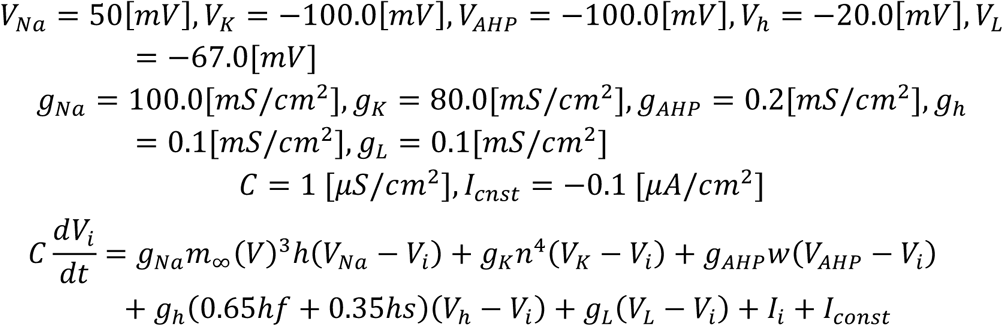

The channel variable h is determined as

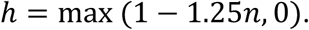

Other channel variables evolve according to

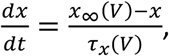

where *x* = *m, n, w, hf* and *hs*. Among these variables, *m* and *n* are determined by

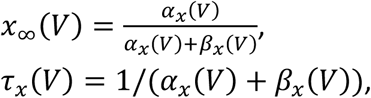

where *x* stands for either *m* or *n*, and

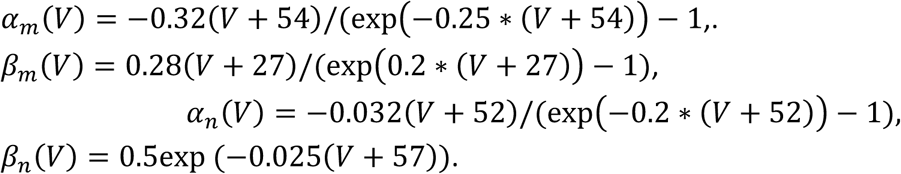

Other variables are determined by

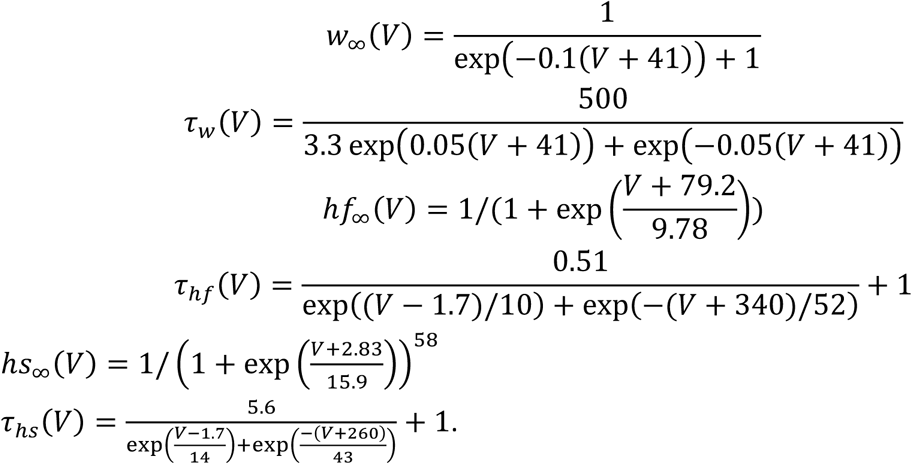

##### b) PV in CA1, MECIII, MECV

We modeled PV neurons in CA1 as well as those in MECIII and MECV as described in (Wang & Buzsáki, 1996).

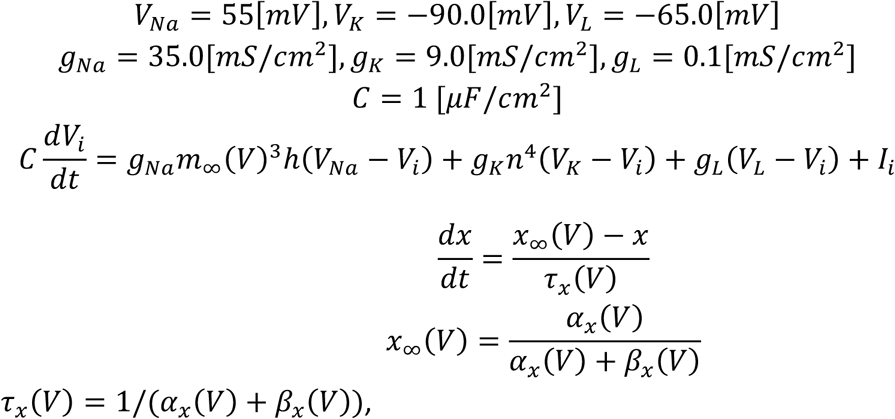

where the index *x*=*n, h* and *m*.

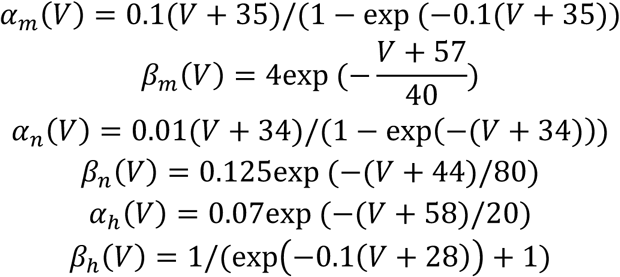

##### c) OLM in CA1

We modeled OLM neurons as described in (Wulff et al., 2009).

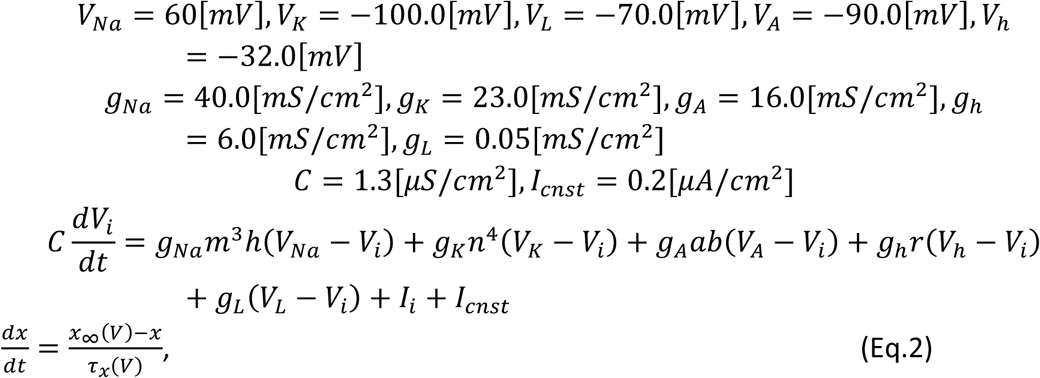

where *x* = *m, h, n, a, b* and *r*. The channel variables are determined by

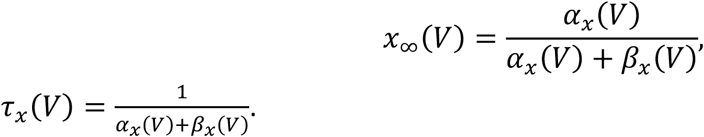

for *m, h* and *n* with

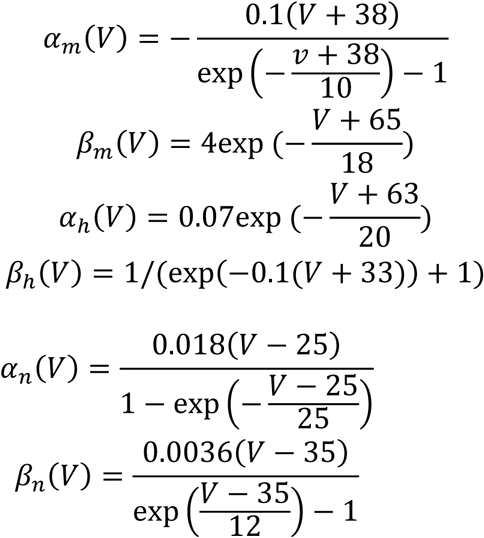

For *x* = *a, b* and *r*, variables in Eq.2 are determined by

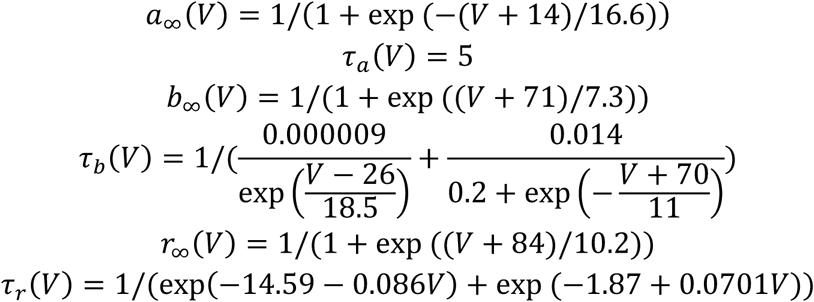

##### d) Excitatory neurons in MECIII

We modeled excitatory neurons in MECIII based on an excitatory neuron model of MECII (S. Middleton et al., 2008), with some modifications of parameter values. These neurons have an AHP current in addition to the standard sodium, potassium and leak currents. The model is described as

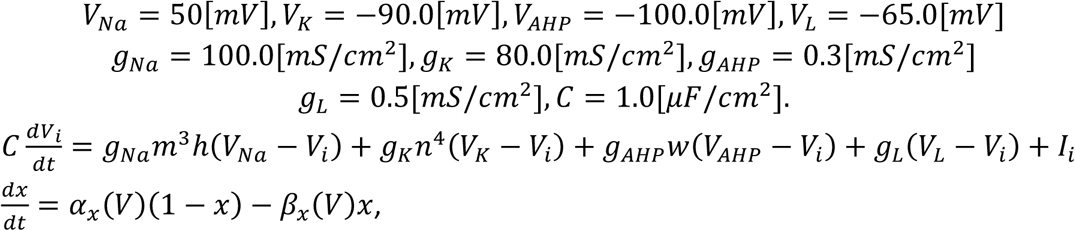

where *x* = *m, n* and *h*.

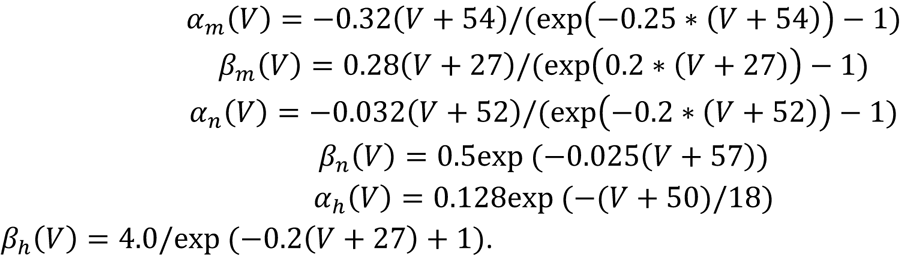

The channel variable *w* is determined by

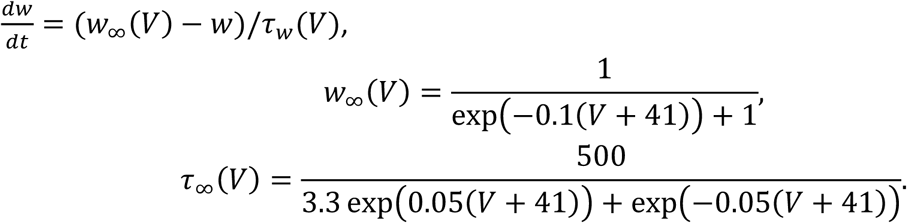

##### e) Excitatory neurons in MECV

Excitatory neurons in MECV are modeled based on (Egorov, Hamam, Fransen, Hasselmo, & Alonso, 2002; Fransén et al., 2006) with simplification of conductance of nonspecific calcium-sensitive cationic, *g*_*CAN*_ (see Disinhibitory system in Materials and methods). Ca flux is regulated through CaL channel.

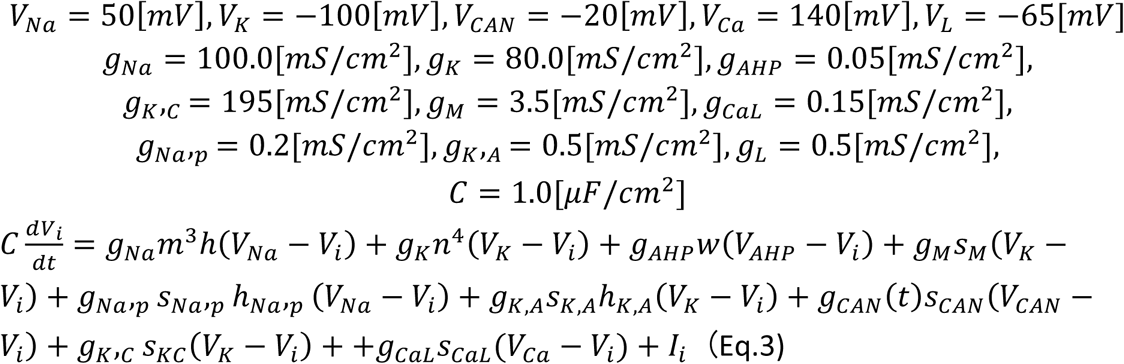

In addition to voltage dynamics, concentration of Ca^2+^, [Ca^2+^], in a neuron is modeled according to:

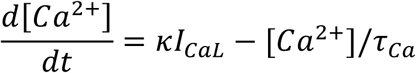

Here, *κ* = 0.5181937[*Fd*^−1^] and *τ*_*Ca*_ = 250[*ms*].

The standard current of sodium and potassium in Eq.3 are exactly same as in excitatory neurons in MECIII. AHP, KA, KC, M, Na_p_ and CaL currents follow the standard activation-inactivation forms:

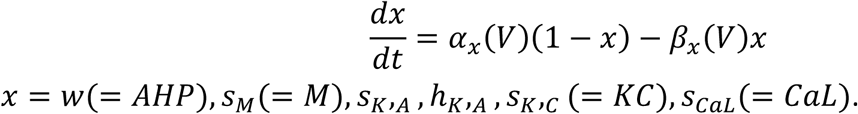

*α* and *β* of each current are according to the following equations.:

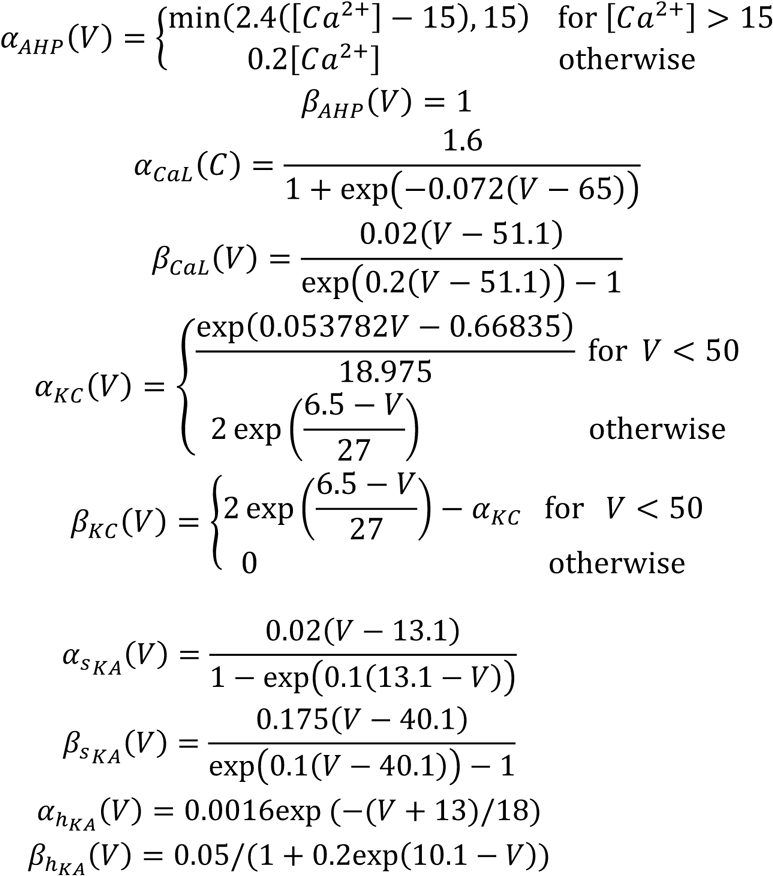

Variables for M, Na_p_ and CAN channels follow the form:

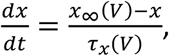

where *x* = *s*_*Na*_,_*p*_, *h*_*Na*_._*p*_, *s*_*M*_(= *M*), *s*_*CAN*_(= *CAN*). *x*_∞_ and *τ*_*x*_ for each variable are according to

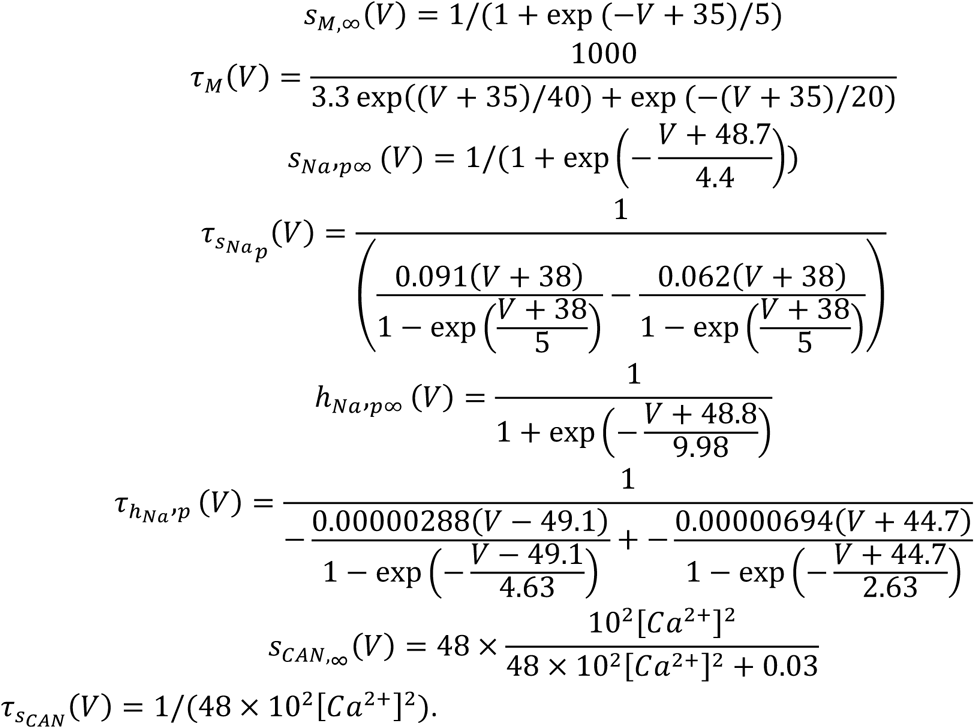

### Synaptic Inputs

Synaptic input *I*_*i*_ to neuron *i* includes excitatory 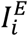 and inhibitory 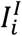 currents. Only in CA1 neurons, OLM currents, which are from OLM neurons, are modeled as another type of inhibitory currents based on (Wulff et al., 2009).

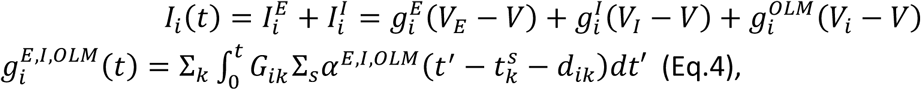

where k is index of pre-synaptic neuron. For the excitatory current, k corresponds to index of excitatory neurons in all cortical areas and excitatory noise neurons projecting to post-synaptic neuron *i*. For the inhibitory currents, *k* corresponds to index of inhibitory neurons (PV, OLM, VIP, GABA) in all cortical areas and inhibitory noise neurons projecting to post-synaptic neuron *i*. Connections *G* among these neurons are described in Circuit model in Materials and Methods. *s* is index of spikes in *k* neuron. *d*_*ik*_ is transduction time lag. When neurons *i* and *k* are within a same area, *d* is chosen randomly from (0,2) ms, while *d* is chosen from (10,15) ms for neurons in different areas. Double exponential functions, *α*, are described as:

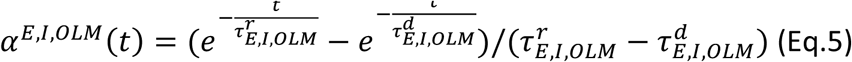

Rise time constant *t*^r^ is 0.05,0.07 and 2.0 ms for E, I, and OLM, respectively, while decay time constant *t*^d^ is 5.3, 9.1 and 22.0 ms for E, I, and OLM, respectively.

### Circuit structure

Our model has three cortical areas (Figure1A) and external oscillating neurons in MECII, CA3 and MS. excitatory neurons in each cortical area are divided into two or four subgroups (Table1). CA1 has four groups denoted as Left (L), Right (R), Center (C) and Home (H). MECIII and MECV have two groups denoted as L and R.

**Table 1:**
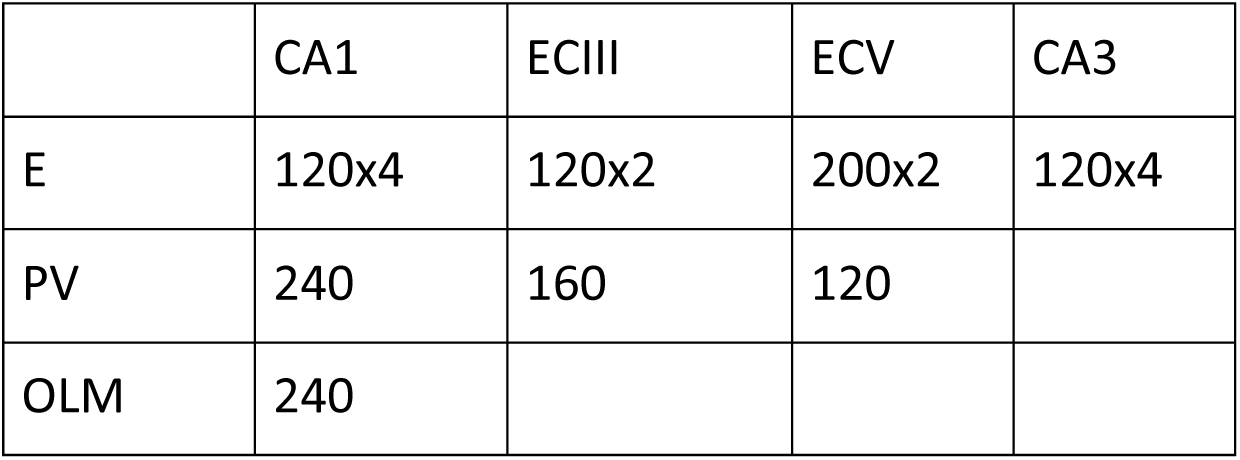
number of neurons in each are.

In addition, a model has 240 VIP neurons in CA1, 120 excitatory neurons in MECII and 360 GABAergic neurons in MS. The GABAergic neurons are divided into three groups projecting to different types of neuron each of which has 120 neurons; groups to OLM, to PV in CA1 and to PV in MECV.

Structure of a circuit is given by a matrix *G* in Eq. 4, which represents efficacy of synaptic connections. Connection between presynaptic neuron *j* and postsynaptic neuron *I, G*_*ij*_ is determined by

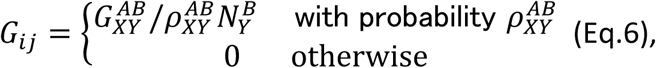

where X and Y are neuron types that *i* and *j* neurons belong to, respectively. A and B refer subgroups (L,R,C,and H) that *i* and *j* neurons belong to, respectively. Because only excitatory neurons in each area are divided as the subgroups, A and B are neglected for other types of neurons. *N*_*Y*_ is the number of type *Y* neurons. If the type Y neurons are divided into the subgroup, *N*^*B*^_Y_ indicates the number of type Y neurons in subgroup B. *G* and *ρ* for each connection are as follows:

#### i) Connection from noise neurons

Connections *G* from noise neurons to the neurons in cortical areas are shown in Table2. A postsynaptic neuron receives inputs from 40 excitatory and 40 inhibitory neurons (*ρ*_*XY*_ = 1, *N*_*XY*_ = 40).

**Table 2.** Connection *G*_*XY*_ from noise neurons to each type of neurons.

| Post-synaptic X<br>Pre-synaptic Y | CA1 |  |  | ECIII |  | ECV |  |
| --- | --- | --- | --- | --- | --- | --- | --- |
|  | E | PV | OLM | E | PV | E | PV |
| Excitatory noise neurons | .32 | .8 | 3.2 | .8 | .2 | .2 | .4 |
| Inhibitory noise neurons | .6 | .2 | .4 | .6 | .8 | .2 | .6 |

#### ii) Connection from excitatory neurons in MECII, MS and CA3

a) connection from excitatory neurons in MECII to PV in MECIII

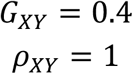

b) from GABAergic neurons in MS

to PV in MECV

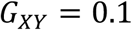

to PV in CA1

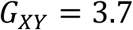

To OLM in CA1

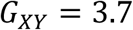

For all connections

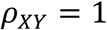

c) from CA3 to E in CA1

for *A* ≠ *B*

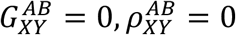

For *A* = *B*

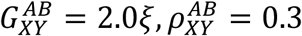

here, *ξ* is chosen randomly from (0,1).

d) from CA3 to PV in CA1

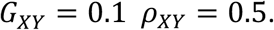

e) from VIP to PV and OLM in CA1

Efficacy of these connections is modified by [ACh] (Tremblay, Lee, & Rudy, 2016) as follows:

For X=PV

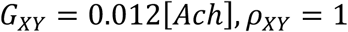

For X=OLM

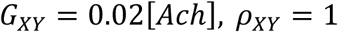

f) Connections between E neurons within the same cortical areas

For *A* ≠ *B*

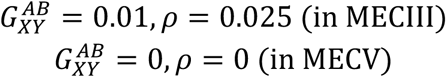

For *A* = *B*

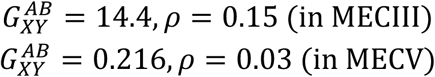

There is no connection between excitatory neurons in CA1 in our model.

g) Connection between neurons within the same cortical areas (except E-E connections) There is no connection between OLMs(Wulff et al., 2009). Connections from OLM to E neurons in CA1 are described in h), since OLM is observed to be attached on proximal dendrite of excitatory neurons in CA1 and regulate inputs from MECIII.

Other connections within cortical areas are shown in Table 3.

**Table 3:** connection parameters in CA1, MECIII, MECV

|  | CA1 |  |  |  |  |  |
| --- | --- | --- | --- | --- | --- | --- |
| (X,Y) | E,PV | PV,E | PV,PV | PV,OLM | OLM,E | OLM,PV |
| $G_{XY}$ | 2.2 | 0.2 | 0.3 | 0.5 | 0.5 | 0.2 |
| $\rho_{XY}$ | 0.5 | 0.5 | 0.5 | 0.5 | 0.3 | 0.5 |

| MECIII |  |  | MECV |  |  |
| --- | --- | --- | --- | --- | --- |
| E,PV | PV,E | PV,PV | E,PV | PV,E | PV,PV |
| 2.5 | 0.5 | 0.01 | 0.2 | 0.3 | 0.01 |
| 0.5 | 0.3 | 0.3 | 0.5 | 0.3 | 0.3 |

h) connections from neurons in CA1 to those in MECV

For X=excitatory neurons in MECV, Y=excitatory neurons in CA1,

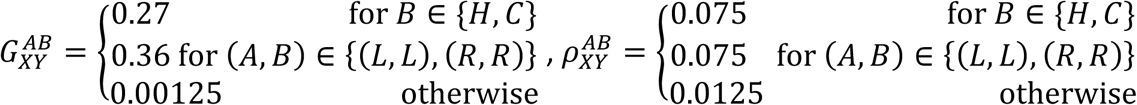

For X=PV neurons in MECV, Y=excitatory neurons in CA1,

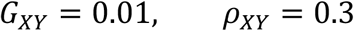

i) connections from neurons in MECV to those in MECIII

For X= excitatory neurons in MECIII, Y=excitatory neurons in MECV,

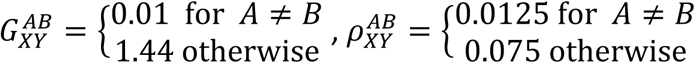

j) connections from neurons in MECIII to those in CA1

For X=excitatory neurons in CA1, Y=excitatory neurons in MECIII,

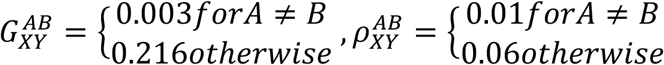

In addition, effect of OLM is implemented for modulating inputs from MECIII as follows: Up to 5 ms after a spike from OLM to E neurons in CA1, efficacy of connections from E neurons in MECIII to this E neurons in CA1 is reduced with multiplication by 0.1.

### Nonlinear interaction of excitatory neurons in CA1 with input from MECIII and CA3

E neurons integrate spikes from CA3 and MECIII (Bittner et al., 2015). We introduced this effect in our model. A prolonged EPSC *I*_*prol*_ is supposed to be generated, when the following conditions are satisfied:

- An excitatory neuron in CA1 receives a burst (three spikes within 15ms) from MECIII and it does not receive any inhibitory input from OLM within 15ms at time t_1_.
- This excitatory CA1 receives a burst (three spikes within 10ms) from CA3 within 20ms from t_1_ (denoted t_2_).
- Previous prolonged EPSC is 100ms earlier than t_2_.

If these conditions are satisfied, a prolonged (100ms) EPSC in E neuron in CA1 is generated according to:

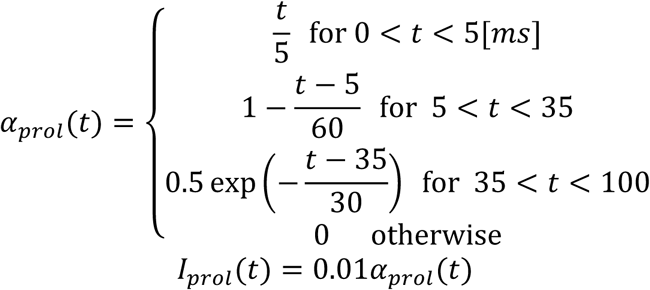

### Disinhibitory system and modulated conductance of nonspecific calcium-sensitive cationic (CAN) current through ACh

We assume that concentration of ACh represents cognitive states and changes dependent of current locations:

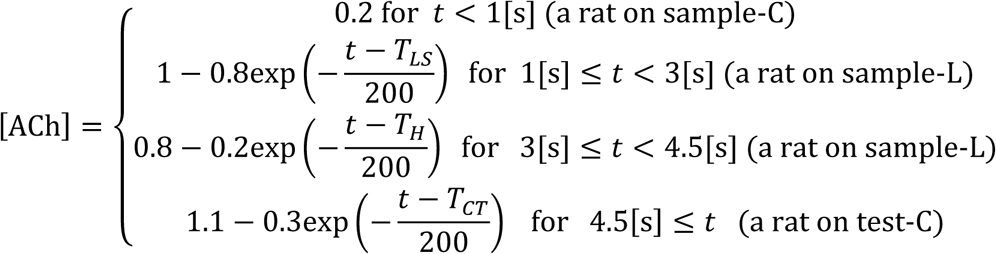

[ACh] modifies neural behavior in a circuit in two pathways. Efficacy of connections from VIP is modified with [ACh] (Circuit structure in Materials and methods) as well as conductance of *I*_*CAN*_. A channel of *I*_*CAN*_ takes high and low conductance states alternatively. Maximum conductance *g*_*CAN*_ is determined by [ACh] and ratio of high conductance state *r*_h_. *r*_h_ is dependent on history of [Ca2+] and [ACh] (Fransén et al., 2006) as follows:

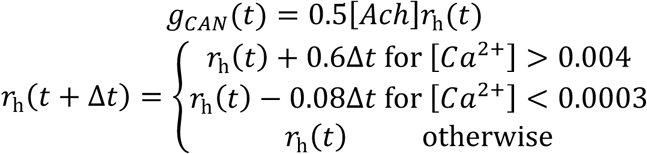

We also set upper and lower bounds for *r*_h_ at 1 and 0.12, respectively.

### Periodicity analysis

Neural activity in the model oscillates driven by external theta rhythm (10Hz). We evaluated how strongly neurons oscillate in the theta rhythm by calculating an autocorrelation function R(t) (Yamamoto et al., 2014). Periodicity index is defined as 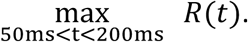

### Experimental data in rodents

Spiking activities and LFP data in CA1 are obtained from previously published data in (Yamamoto et al., 2014) for Figures 6B and 7C and that in (Mizuseki et al., 2013) for Figures 6D and 6E. Experiments were approved by the Institutional Animal Care and Use Committee of Rutgers University. All procedures for animal care and use were performed in accordance with the National Institutes of Health Guide for the Care and Use of Laboratory Animals. Detailed conditions on data recording are described in these papers.

LFP activities were band-pass filtered (6-12Hz) as theta wave and instantaneous phase of the filtered theta wave were derived from Hilbert transform. For spiking activities, we dropped a part of spikes to analyzed as follows:

Spiking activities in (Yamamoto et al., 2014) were recorded with the silicon linear probes and were analyzed as multi-unit activities. In this paper, however, we roughly distinguish putative excitatory neurons from inhibitory ones in order to show clear theta preference. According to (Mizuseki et al., 2009), we sorted spikes by trough to peak latency. Due to short length of single spike profile in the data, we cannot identify baseline before spike and consequently cannot compute peak amplitude asymmetry. We assigned spikes with the latency larger than 0.5ms to putative excitatory neurons. For Figure 6B, we have the sorted spikes in three out of five rats because spikes of the rest two rats do not show clear theta preference due to small number of spikes.

For spiking activities in (Mizuseki et al., 2013), we used sessions “ec014.12”,”ec014.16”,”ec014.17”,”ec014.27”,”ec014.28”,”ec013.44”,”ec013.46”,”ec0 16.30”. Because phase shift in the side arm was observed only in deeper neurons in (Fernández-Ruiz et al., 2017), we only used spikes of deeper neurons according to (Mizuseki, Diba, Pastalkova, & Buzsáki, 2011).

### Calculations of the ratio of spikes at the peaks in theta and the side-to-center ratio

In Figure 6F, we used the same neurons in Figure 6E (see Experimental data in rodents). Further, we filtered these neurons by two criteria: i) number of spikes in the later center arm are larger than 50; ii) the ratio of spikes in the later center arm to total spikes is larger than 5%. To exclude place cells representing the earlier center arm and junction of T-maze (since these cells are likely to spike at the peaks in theta oscillation in the later center arm), we further excluded the cells that spike in the later center arm less frequently than at the earlier center arm or at the junction. Eleven neurons remained. For these neurons, we calculated the ratio in number of spikes around the peaks (90-270 degree) to all spikes in the later center arm. We call this quantity the ratio of spikes at the peaks. We also calculated the ratio of average number of spikes emitted in the later center arm to that of average number of spikes emitted in the side arms. We call this quantity the side-to-center ratio.

### Data and Software Availability

The computer codes used to generate the present simulation results will be available upon request.

## Acknowledgement

We thank Jun Yamamoto for providing experimental data and fruitful discussion and György Buzsáki for publicly sharing the valuable data. This work was supported by KAKENHI (nos. 18H05213, 18K15343 and 19H04994) from the MEXT, Japan.

